# Discovery of Tcf7 regulators with clonally-resolved CRISPR screens identifies Trim28 as a mediator of CD8 T cell differentiation in tumors

**DOI:** 10.64898/2026.06.14.732162

**Authors:** Marc A. Schwartz, Dillon Lue, Mohammed Mutaher, Miriam McNerney, Oluwatomisin Olajide, Arnav Mehta, Jared H. Rowe, Ryan J. Park, Mohsen Malehmir, Juan Dubrot, Aziz M. Al’Khafaji, Matthew Bakalar, Lynn Bi, Martin W. LaFleur, Kathleen B. Yates, Robert T. Manguso, John G. Doench, Arlene H. Sharpe, Yuri Pritykin, Nir Hacohen

**Affiliations:** Department of Pediatric Oncology, Dana-Farber Cancer Institute, Boston, MA, USA; Division of Hematology/Oncology, Boston Children’s Hospital, Boston, MA, USA; Broad Institute of MIT and Harvard, Cambridge, MA, USA; Department of Computer Science, Princeton University, Princeton, NJ, USA; Lewis-Sigler Institute for Integrative Genomics, Princeton University, Princeton, NJ, USA; UMass Chan Medical School, Worcester, MA, USA; Massachusetts General Hospital Cancer Center, Boston, MA, USA; Harvard Medical School, Boston, MA, USA; Division of Radiation Oncology, MD Anderson Cancer Center, Houston, TX, USA; Solid Tumors Program, Cima-Cancer Center Clínica Universidad de Navarra (CCUN), Pamplona, Spain; Broad Clinical Labs, Cambridge, MA, USA; Department of Immunology, Blavatnik Institute, Harvard Medical School, Boston, MA, USA; Gene Lay Institute of Immunology and Inflammation, Brigham and Women’s Hospital, Massachusetts General Hospital, and Harvard Medical School, Boston, MA, USA; Krantz Family Center for Cancer Research, Massachusetts General Hospital, Boston, MA, USA; Department of Medicine, Massachusetts General Hospital, Harvard Medical School, Boston, MA, USA

## Abstract

Stem-like TCF7+ CD8 T cells sustain anti-tumor responses and support immune checkpoint blockade. We systematically identified regulators of this cell state using genome-wide CRISPR screens in primary T cells *in vitro*. Using random barcodes to link clonal relationships with guide identity and transcriptional states in single cells, we inferred differentiation trajectories and differentiation rates of CD8 T cells in tumors, while mitigating confounding clonal bias. We found that Trim28-deficient T cells in tumors were enriched in the TCF7+ cell state, depleted in cycling and terminal effector states, and uniquely generated a tissue-resident memory (TRM)-like state with increased chromatin accessibility at known TRM loci as well as repeat elements. Despite the increase in TCF7+ CD8 T cells, loss of Trim28 did not improve tumor control, likely because of reduced effector differentiation, highlighting the need for tuning the balance and dynamics of stem-like versus effector states for effective tumor clearance.

## Introduction

A central challenge in cancer immunotherapy is to identify the CD8 T cell states that sustain durable tumor control and determine how those states can be therapeutically shaped. Recent studies in mouse models and human tumors have highlighted a critical role for a low abundance, stem-like subset of CD8⁺ T cells in sustaining anti-tumor immunity and response to immune checkpoint blockade (ICB).^1–3^ These cells, often characterized by high expression of the transcription factor Tcf7/TCF7 (also known as TCF-1), serve as a self-renewing pool capable of generating more differentiated cytotoxic progeny, coupling long-term persistence with continued effector output.^4,5^ Their presence within the tumor microenvironment has been associated with improved responses to ICB.^6,7^ Under chronic antigen exposure, PD-1⁺TCF1⁺ stem-like exhausted cells provide an anti-PD-1-responsive resource pool, whereas terminally exhausted cells are more cytotoxic, have reduced proliferative potential, and respond poorly to checkpoint blockade.^2–4,8^ Importantly, these TCF7⁺ stem-like cells are not only predictive of therapeutic response but are required for the generation of long-lasting immunity, as they retain proliferative capacity and the potential to repopulate the effector T cell compartment. This also creates a key therapeutic tradeoff: durable immunity requires preservation of a renewable stem-like pool, while tumor clearance requires sufficient generation of differentiated effector cells.

Despite their therapeutic relevance, relatively little is known about the upstream pathways that regulate Tcf7 expression and the establishment of the stem-like T cell fate. While the transcriptional and epigenetic profiles of Tcf7⁺ cells have been characterized in both viral and tumor settings, the signals that govern whether an activated CD8⁺ T cell adopts a stem-like versus terminally differentiated state remain incompletely understood. Furthermore, the tumor microenvironment imposes unique metabolic, inflammatory, and antigenic stresses that may constrain the generation or maintenance of stem-like T cells, presenting both a challenge and an opportunity for therapeutic modulation. Current immunotherapy strategies have largely focused on enhancing effector T cell activity or preventing terminal exhaustion, but there are no approved therapies specifically designed to promote the formation or expansion of Tcf7⁺ stem-like CD8⁺ T cells. Identifying genetic regulators of Tcf7 expression could thus yield new targets for interventions that increase the stem-like T cell pool, improve T cell persistence, and ultimately enhance the efficacy of ICB and other immunotherapeutic approaches.

Prior *in vivo* single-cell CRISPR screening has mapped transcriptional circuits that control intratumoral CD8⁺ T cell differentiation and proliferation, including the IKAROS-TCF-1, ETS1-BATF and RBPJ-IRF1 axes.^9^ Two limitations of this approach are first that activation before adoptive transfer can alter T cell state before tumor encounter, and second, without clonal barcodes, stochastic expansion of individual T cell clones cannot be distinguished from perturbation-specific effects on cell number or cell-state composition.

Several studies have used clonal information to learn how individual cells diversify during differentiation. In infection models, lineage tracing showed that individual naive CD8⁺ T cells bearing the same T cell receptor (TCR) can generate heterogeneous expansion and differentiation patterns.^10^ In hematopoiesis, cellular barcoding showed that early progenitors can have diverse and heritable lineage biases^11^, and subsequent lineage tracing with single-cell RNA-seq linked early transcriptional state to later clonal fate.^12^ A recent study used barcodes in bulk to improve statistical power in CRISPR screens by treating clones as replicates, and to track global versus tissue-level clone expansion^13^, while another study in cortex development *in vitro* used barcodes to determine lineage relationships in the context of perturbation.^14^

Here, we developed a naive-state-preserving lentiviral guide-delivery strategy and combined it with clonally resolved single cell RNA-seq to identify regulators of Tcf7 expression and CD8 T cell fate in tumors. This approach nominates Trim28 as a regulator of progenitor and TRM-like CD8 T cell states and provides a framework for separating perturbation effects on clonal expansion, differentiation, and cell-state composition in vivo.

## Results

### A method for transduction of naive CD8 T cells

Methods for lentiviral transduction of CD8 T cells that require TCR activation prior to transduction greatly affect T cell state and result in downregulation of Tcf7, complicating their use in CRISPR screens to study regulation of Tcf7. To identify key modulators of Tcf7, we developed and validated a direct, naive-state-preserving lentiviral transduction method for delivering CRISPR guide RNA libraries to naive CD8 T cells without TCR activation (Figure 1a). Key features of this approach include mixed VSV-G/MMLV ecotropic lentiviral envelopes, RetroNectin-mediated virus loading, LentiBOOST transduction enhancer, Thy1.1-based enrichment of transduced cells, and culture without TCR stimulation. We reported an IL-7 only version of this platform in prior in vivo CD8 T cell perturbation studies.^13,15^ In the present study, we used IL-7 and IL-15 during transduction to maximize survival and cell recovery while scaling to pooled genome-wide CRISPR screens, motivated by prior work showing that IL-15 can promote survival of naïve CD8 T cells.^16^

**Figure 1:**
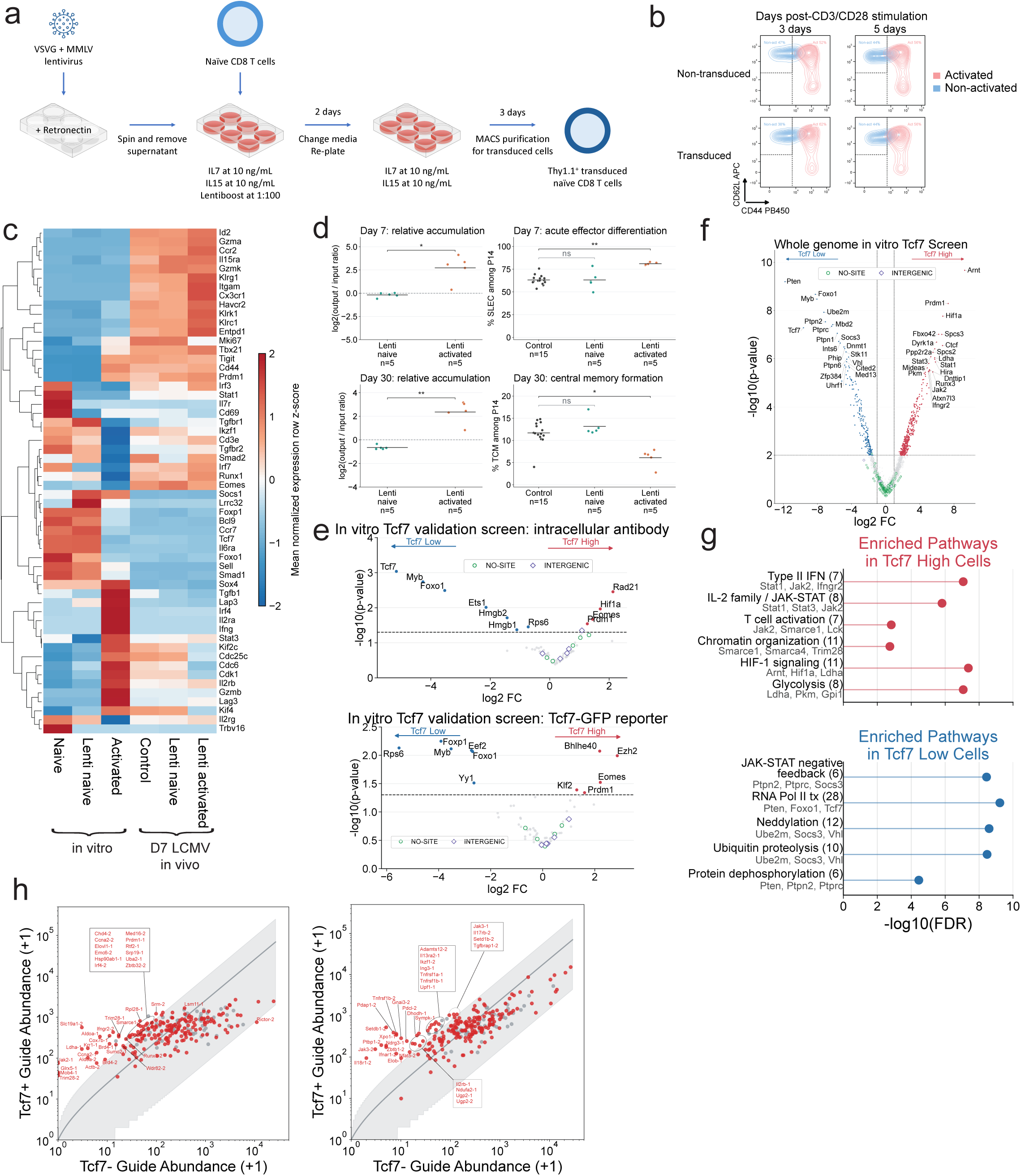
Lentiviral transduction without T cell activation preserves naive state and enables CRISPR screens for regulators of Tcf7 expression. (a) Method for lentiviral transduction of naive CD8 T cells (b) Flow cytometry plots of T cell activation markers after lentiviral transduction and CD3/CD28 stimulation. (c) Heatmap of DESeq2-normalized bulk RNA-seq expression for T cell activation and differentiation genes across pre-transfer P14 CD8 T cell inputs and day-7 cells recovered after LCMV Armstrong infection. Pre-transfer samples include freshly isolated naive cells, lentivirally transduced naive cells using our approach, and anti-CD3/CD28-activated non-transduced cells. Day-7 samples include cells that were derived from cotransferred control cells with lentivirally transduced naive cells using our approach or lenti-activated cells recovered from spleens of infected recipient mice. (d) Competitive transfer of congenically marked P14 LCMV-specific CD8 T cells where freshly isolated naive control P14 cells were mixed with naive lentivirally transduced or activated transduced P14 cells. Left, change in the comparator-to-control P14 cell ratio between cells prior to transfer (input) and cells after recovery (output) at days 7 and 30 after LCMV Armstrong infection. Right, flow-cytometric quantification of the percentage of recovered P14 cells in each transferred group with a short-lived effector cell (SLEC; KLRG1+CD127-) phenotype at day 7 or central memory T cell (TCM; CD62L+CD44+) phenotype at day 30. n is the number of recipient mice. P-value annotations were calculated with two-sided Welch t tests comparing naive transduced and activated pre-transfer cells for ratio changes and two-sided paired t tests comparing each experimental population with matched control cells for SLEC and TCM phenotypes. *, Benjamini-Hochberg adjusted P < 0.05; **, Benjamini-Hochberg adjusted P < 0.01 (Table S1.1). (e) Volcano plots quantifying results of *in vitro* CRISPR screens using a 70-gene gRNA library enriched for likely regulators of Tcf7. Cell populations were sorted using either intracellular Tcf7 antibody staining or a Tcf7-GFP intronic reporter. Target genes with P <= 0.05 were highlighted (Table S1.2; Table S1.3). (f) Volcano plot quantifying results of *in vitro* whole genome CRISPR screen. The top 20 most significant genes in both directions were labeled (Table S1.4). (g) Pathway enrichment summary for significant genes from the whole-genome *in vitro* Tcf7 CRISPR screen. (h) Scatter plots of Tcf7+ guide abundance (y-axis, log scale) versus Tcf7- guide abundance (x-axis, log scale), after adding a pseudocount of 1 to value on both axes, for two *in vivo* Tcf7 sort screens. Control guides are plotted in gray and gene-targeting guides in red. Gray line and shaded region represent the modeled relationship and 95% predictive interval relating Tcf7-guide abundance to Tcf7+ guide abundance, fit using control guides (Methods). Gene-targeting guides with nominal P < 0.05 were labeled (Table S1.6; Table S1.7).

We validated this method by determining whether transduced CD8 T cells remain phenotypically naive in culture and behave similarly to freshly isolated naive CD8 T cells in the context of a subsequent *in vivo* LCMV infection model. Following transduction with this method, we found that transduced naive CD8 cells retained a CD62L-high CD44-low naive phenotype, and acquired the same activation phenotype as untransduced cells after CD3/CD28 stimulation *in vitro* (Figure 1b).

To test the ability of naive transduced CD8 T cells to respond to an antigen and form functional memory cells and precursors in mice, we used a lymphocytic choriomeningitis virus (LCMV) model. Competitive transfer experiments were performed using: (1) P14 LCMV-specific CD8 T cells transduced with our naive state-preserving lentiviral transduction method (termed lenti naive); (2) P14 cells transduced and activated with anti-CD3/CD28 stimulation (termed activated condition). Both transfers were mixed with freshly isolated naive P14 CD45.1+CD45.2+ CD8 T cells (termed control cells). Recipient mice were then infected with LCMV Armstrong, and splenic T cells were analyzed 7 and 30 days following infection. Compared with activated cells, lenti naive cells showed a transcriptomic profile more similar to naive cells before transfer and at day 7 after LCMV infection (Figure 1c; Figure S1a-c). Flow cytometry showed that T cell abundance was comparable for transduced and untransduced naive T cells, but transduced activated T cells were substantially higher in abundance (Figure 1d). In addition, activated transduced T cells led to increase in short lived effector cells at day 7 and decrease in proportion of central memory and memory precursor populations at day 30, but there were no differences in these populations after transfer of transduced naive cells relative to control (Figure 1d). Furthermore, we also flow-sorted central memory and effector memory P14 CD8 T cells at day 30 and then performed RNA-seq. Central memory and effector memory cells derived from lenti-naive P14 cells more closely resembled the corresponding memory populations from cotransferred naive controls than did memory cells derived from lenti-activated P14 cells (Figure S1d). In summary, naive CD8 T cells transduced using our method behave similarly to freshly isolated naive T cells in their abundance and cell state distribution *in vivo*.

### CRISPR screens to identify regulators of Tcf7 and other transcription factors

In order to evaluate if we could perform CRISPR screens, we tested libraries of known positive and negative regulators of Tcf7 *in vitro*. These libraries were transduced into naive CD8 T cells, isolated 5 days post-transduction to allow time for gene deletion, and then activated using anti-CD3/CD28 antibodies to evaluate which gene deletions modulate activation-induced downregulation of Tcf7 expression. One week post-anti-CD3/CD28 stimulation, cells were either fixed and stained intracellularly for Tcf7 protein, or monitored as live cells using a Tcf7-GFP reporter. Cells were sorted into low or high Tcf7 expression bins, and gRNA abundance was determined per bin by sequencing. Known regulators of Tcf7 scored in these pilot screens with good separation between the top hits and control gRNAs, and these results were largely consistent (Myb, Foxo1, Prdm1, Eomes) between the intracellular staining and reporter conditions, except the Tcf7 locus continues to display reporter expression even when Tcf7 gene is targeted as well as other genes that may show differences via unknown mechanisms (Figure 1e).

To identify novel regulators of Tcf7 in CD8 T cells, we used the same pipeline as above using a genome-wide CRISPR library in vitro (Figure 1f), and found the same known regulators as in the pilot tests. Pathway enrichment of Tcf7-high and Tcf7-low hits highlighted broad programs related to T cell activation and differentiation, chromatin organization, and metabolic regulation (Figure 1g).

Additional genome-wide screens across Prdm1, Eomes, Tox, and Tcf7 stimulation contexts revealed distinct pathway-enrichment patterns for each transcription factor (TF), but repeatedly implicated broad programs associated with T cell receptor/cytokine signaling, PIP3-AKT/mTOR/HIF-1 signaling, chromatin and transcriptional regulation, cell cycle, DNA replication, and rRNA processing (Figure S2a,b; Table S1.5).

To identify the subset of hits from our *in vitro* Tcf7 screen that also function *in vivo*, we adoptively transferred naive Cas9+ CD8 T cells transduced with gRNA libraries targeting 100 genes with the strongest effect on Tcf7 expression *in vitro*. T cells were isolated from implanted B16-OVA tumors ∼2 weeks post transfer, and sorted for Tcf7-GFP reporter high and low populations to determine guide enrichment or depletion (Figure 1h). We nominated ten genes based on strong phenotypes and potential for discovering new mechanisms.

### Gene deletion in tumors alters T cell state distributions and reveals TRM-like cells after Trim28 or Setdb1 loss

To better understand the roles of these genes in T cell differentiation, we designed a pool of gRNAs with two gRNAs (selected for strongest efficacy from prior screen) for each of the 10 genes, together with no-site and intergenic control gRNAs. Naive T cells were transduced with the pool and transferred into B16-OVA tumor-bearing mice. On day 10 after T cell transfer, mice were treated with anti-PD-1 antibody or isotype control for 5 days. T cells were harvested and donor cells sorted with Thy1.1, and profiled using droplet-based scRNA-seq for cellular mRNAs, gRNAs (with linked unique molecular identifiers [UMIs]), and unique antibody-based hashtags per mouse (Figure 2a). Guide RNA assignment was detected for cells; cells assigned only to control gRNAs are referred to as NT controls; cells without a detected gRNA assignment are referred to as no-guide (NG) cells.

**Figure 2:**
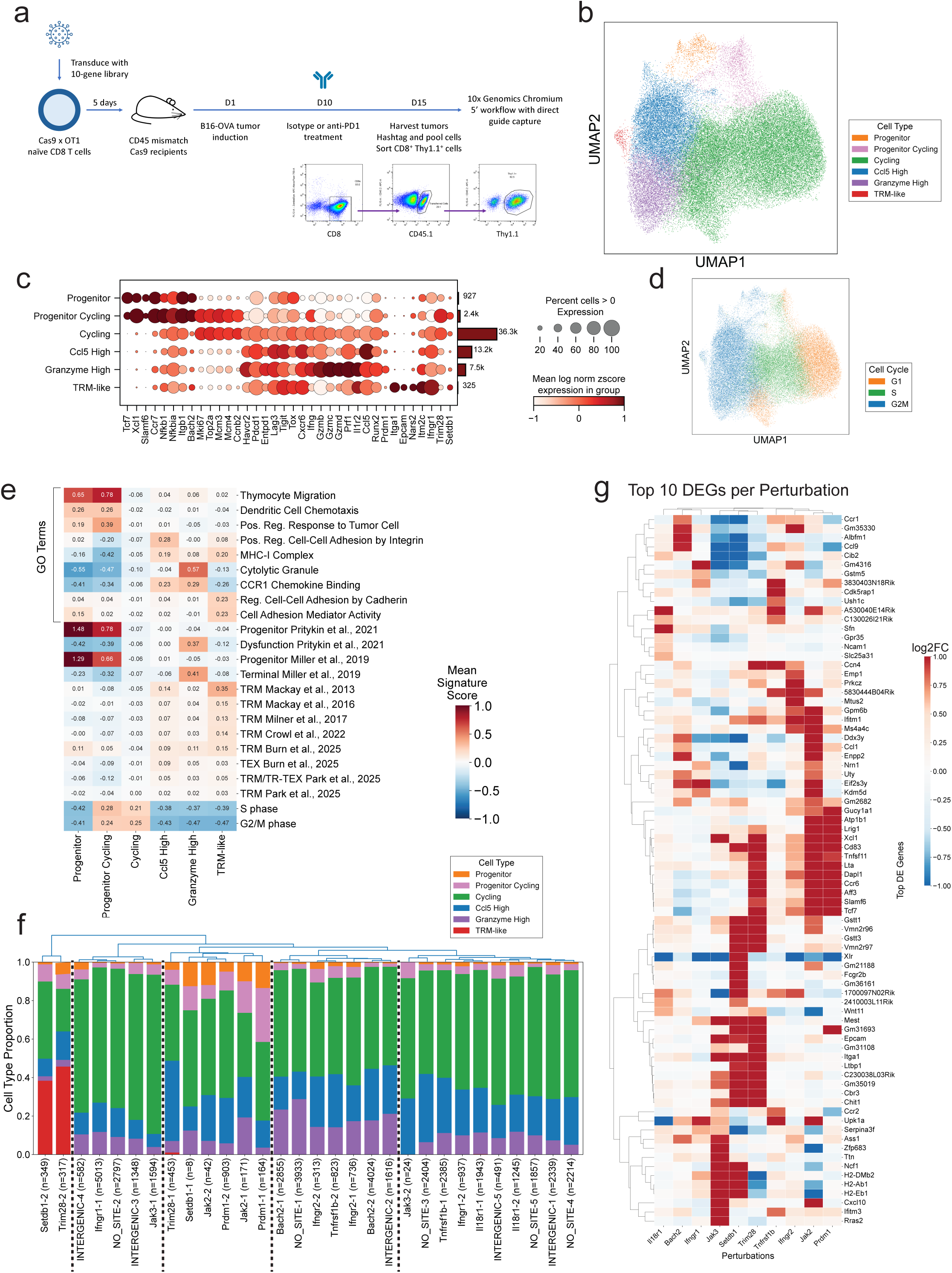
*In vivo* Perturb-seq of CD8+ tumor-infiltrating lymphocytes (TILs) reveals cell type distribution changes. (a) Schematic of the experimental design for clonal Perturb-seq involving naive Cas9 x OT1 CD8+ T cells transduced with the 10-gene Perturb-seq library, which contained 2 gRNAs per gene, 5 no-site control gRNAs, and 5 intergenic control gRNAs. Following transduction, cells were adoptively transferred into mice. Tumor injection occurred 1 day later, followed by isotype or anti-PD-1 treatment on day 10. Transferred cells were harvested from the tumor and processed for single-cell RNA-seq after 5 days. (b) UMAP representation illustrating cell state annotation from clusters obtained by Leiden clustering of the kNN graph of cell-cell transcriptomic similarities. (c) Dot plot visualizing the expression levels of key markers across identified cell states across all cells. Dot size represents the percentage of cells expressing each marker, while color intensity shows z-score normalized log expression levels. The bar plot on the right of the dot plot shows the number of cells per cell type. (d) UMAP plot displaying cell cycle phases. (e) Heatmap of mean signature scores across CD8+ TIL states for: selected GO terms; progenitor/dysfunction^60^; progenitor vs terminal exhaustion^2^; TRM signatures from acute infection comparing tissue-resident vs circulating memory after clearance^17–20^; TEX versus TRM signatures defined from CD8+ T cells in tumor versus paired non-cancerous tissue^21^; TRM (acute, Arm) and TR-TEX (chronic, Cl13)^22^; and cell-cycle (S phase, G2/M). Numeric labels denote the mean signature score per cell type. Representative genes underlying the GO, progenitor/exhaustion, and TRM/TEX signature programs are shown (Figure S3c-e). (f) Stacked bar chart depicting the proportion of each cell state within groups targeted by specific gRNAs; x-axis labels include n = number of cells per guide. (g) Heatmap showcasing the top ten upregulated differentially expressed genes (DEGs) with log2FC > 0.5 for each perturbation where the comparison is cells with the perturbation vs non-targeting cells.

Unsupervised clustering of CD8⁺ TILs resolved six states (Figure 2b; Figure S3a): two Tcf7-high progenitor populations—one quiescent and one cycling—each enriched for progenitor signatures (Figure 2c,e; Figure S3d); three differentiated populations (Granzyme High with cytolytic-granule and dysfunction/exhaustion programs, Ccl5 High, and a proliferative “Cycling” group); and a novel cluster determined to be TRM-like (Figure 2c,e). The Cycling group could be further clustered by stage-specific cell-cycle genes (G1/S and G2/M), and we did not observe Granzyme High or Ccl5 High subsets within the cycling population (Figure S3b). Differentially expressed genes (DEGs) in progenitor and progenitor-cycling subsets were enriched for migration/chemotaxis terms, consistent with trafficking from lymph node to tumor (Figure 2e; Figure S3c). Ccl5 High and Granzyme High cells showed enrichment for MHC-I complexes, cytolytic granules, and chemokine binding. The TRM-like population was enriched for cell-adhesion processes, consistent with a potential tissue-resident phenotype (Figure S3c).

We further confirmed the annotation of the novel cluster as a TRM-like cell type based on high expression of Itga1 (CD49a), a canonical TRM marker, and its high enrichment relative to other cells in our dataset for multiple published TRM programs (Figure S3e). These include signatures from acute infection models that compare tissue-resident memory to circulating memory after viral clearance^17–20^ and a pan-cancer T cell exhaustion (TEX) versus TRM signature defined from CD8+ T cells in tumor and paired non-cancerous tissue.^21^ Notably, the TRM-like cells from our study do not score as highly as our Progenitors for a distinct TRM program^22^, possibly because this signature captures genes that are more highly expressed post acute infection versus chronic infection. Thus this signature highlights more stem/memory-like T cells which leads to higher expression in the progenitor cells we identified.

We next clustered genetic perturbations based on the proportions of the six CD8+ T cell states. We observed two clusters with distinct phenotypes and lacking control guides: the most distinct cluster with Setdb1 and Trim28 perturbations was defined by the presence of the novel TRM-like state and a higher proportion of progenitors; the second with Prdm1 and Jak2 perturbations by a higher proportion of progenitors. In addition, while perhaps a weaker effect due to the presence of 2 of 10 control guides, a third cluster, which includes Bach2, showed higher numbers of Ccl5 High and Granzyme High (Figure 2f). The remainder of the perturbations clustered with control guides.

Next, we aimed to find shared and distinct transcriptional effects across perturbations based on differential expression in perturbed versus NT cells. We clustered fold-change values for the ten significantly differentially expressed genes with the highest positive log2 fold-change in each perturbation (Figure 2g). Jak2 and Prdm1 perturbed cells clustered together based on progenitor-associated genes (Tcf7, Sell, Xcl1), consistent with the observed higher proportion of progenitors (Figure 2f), while Setdb1 and Trim28 clustered together as a result of forming the TRM-like state, and Bach2 showed the lowest expression of Tcf7 consistent with an increase in differentiated T cells. These Bach2 and Prdm1 phenotypes align with prior work in chronic infection and tumors showing that Bach2 enforces stem-like CD8 T cell programs, whereas Prdm1/Blimp-1 promotes terminal exhaustion programs.^23–25^ Combining the findings from differential expression with cell state proportion changes, we classified perturbed genes into those required for exiting the progenitor state and entering activated states (Prdm1, Jak2), those inhibiting entry into the activated state (Bach2), and those that inhibit the TRM-like state (Trim28, Setdb1).

### Clonal tracking using barcodes reveals clonal heterogeneity

In response to tumor challenge, CD8+ T cells undergo clonal expansion, an essential step for effective anti-tumor immunity. Since prior lineage-tracing studies showed that individual naive CD8+ T cells bearing identical TCRs can generate heterogeneous expansion and differentiation patterns^10^, we developed a clonal barcoding strategy (Figure 3a) to investigate how individual clones respond to genetic perturbations. Naive Cas9-expressing CD8+ T cells were transduced with lentiviruses expressing a gRNA along with a random 6-nucleotide barcode, enabling each individual transduced cell to have a unique guide-barcode combination. After cells divide in response to the tumor antigen, cells with the same gRNA-barcode are identified as a clone derived from a common cell of origin. To assign a unique clonal barcode to a single cell, we developed new approaches for assigning guides and barcodes to cells and calling doublets (Methods; Figure S4). We observed heterogeneity in both clone size and cell-state composition in NT clones (Figure 3b) and clones from a single NT guide such as NO-SITE-4–targeted clones (Figure 3c; Figure S5a).

**Figure 3:**
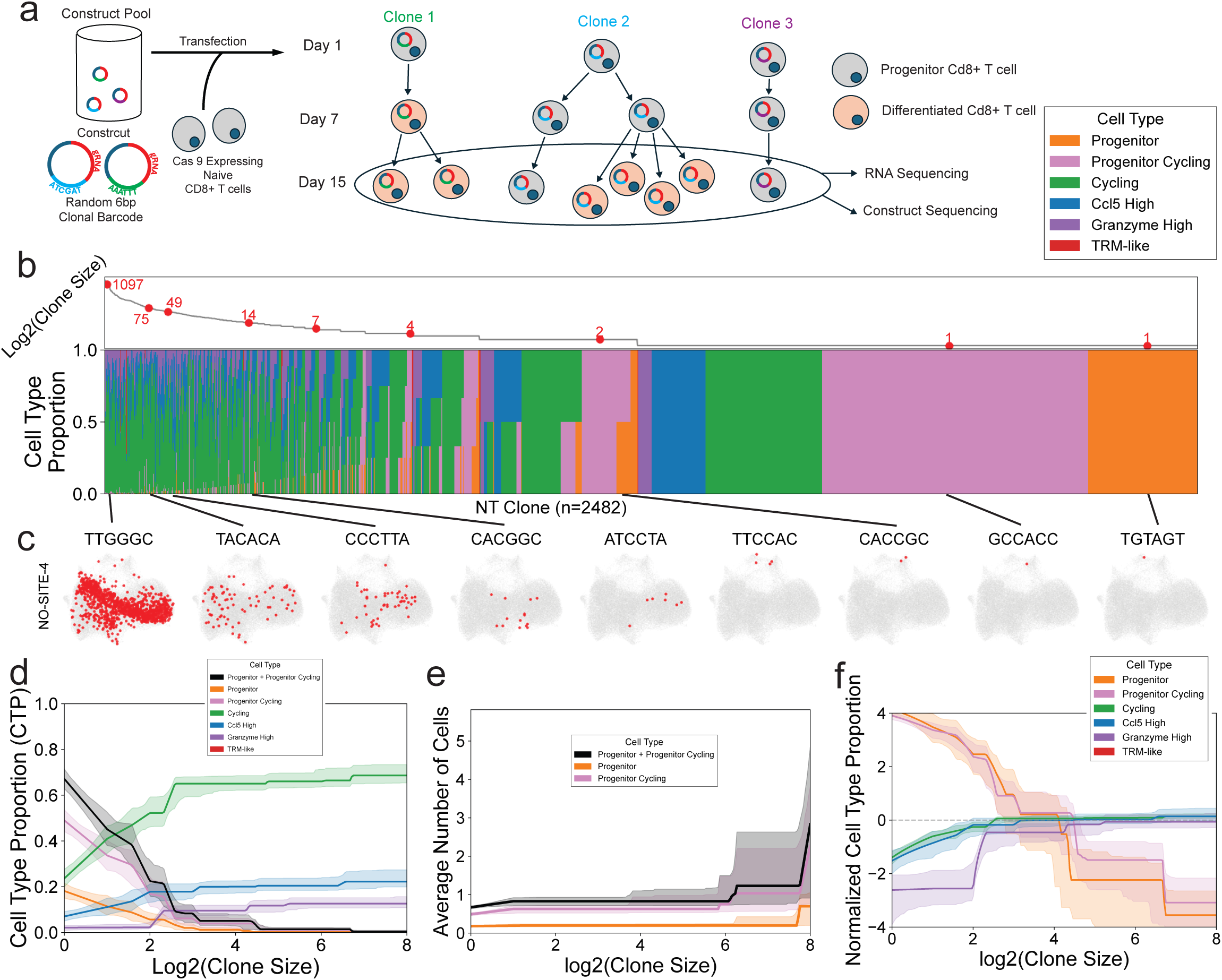
Clonal barcoded lineage tracing of Perturb-seq data reveals clonal heterogeneity. (a) Schematic of the novel clonal barcoding strategy. Naive Cas9-expressing CD8+ T cells were transduced with constructs containing both a unique gRNA and a clonal barcode (colored circles). After adoptive transfer into tumor-bearing mice, cells expanded *in vivo*. Single-cell RNA-seq coupled with construct sequencing to measure the transcriptome for each cell with an assigned gRNA and clonal barcode. (b) Clones ordered by log2(clone size), then by cell-type composition, along the x-axis. Top panel shows log2(clone size) across all clones; red dots mark clones selected from the NO-SITE-4 gRNA set via uniform sampling weighted by log2(clone size) (larger clones are more likely to be selected). Bottom panel shows stacked cell-type proportions for each clone. (c) UMAP visualizations for the same sampled NO-SITE-4 clones, highlighting each clone’s cells in red over all cells (gray). (d) Isotonic regression fits (using NT clones only) showing the relationship between log2(clone size) and the proportion of cells in each state; shaded ribbons indicate 95% bootstrap confidence intervals. (e) Isotonic regression fits (using NT clones only) showing the relationship between log2(clone size) and the number of cells in progenitor and progenitor cycling states; shaded ribbons indicate 95% bootstrap confidence intervals. (f) Normalized isotonic regression fits (using NT clones only) showing the relative enrichment or depletion of each T cell state as a function of clone size. Values represent the log2 ratio of each cell type’s proportion within clones of a given size to its global proportion across all clones; shaded ribbons indicate 95% bootstrap confidence intervals.

To determine quantitatively how cell states vary with clone size for NT clones, we used isotonic regression, a non-parametric method to fit a monotonic trend as a function of log2(clone size) with uncertainty quantified with 95% bootstrap confidence intervals. Although the proportion of differentiated states increases and the proportion of progenitors (i.e., Progenitor and Progenitor Cycling states) drops with increasing clone size (Figure 3d), the progenitor populations remain approximately constant (Figure 3e). We find that Progenitor cells dominate the smallest clones, while Granzyme High and other differentiated states become more prevalent in larger clones (Figure 3e). Clone sizes were highly skewed, with many small clones and relatively few large clones (Figure 3b; Figure S6a). Together these findings are consistent with rapid expansion of differentiated cells.

To compare cell states more directly, we normalized each cell type’s frequency within a given clone size by its global frequency across all clones and plotted the log2 enrichment or depletion (Figure 3f). Treating clone size as a proxy for degree of differentiation, the ordering of the lines in Figure 3f reveals a clear hierarchy: progenitor cells show the strongest enrichment in small clones and the strongest depletion in large clones, followed by progenitor cycling, Ccl5 High, and finally Granzyme High, which becomes depleted at smaller clone sizes but enriched at larger clone sizes. This relative positioning across clone sizes quantitatively defines the differentiation trajectory: Progenitor, Progenitor Cycling, Ccl5 High, and Granzyme High, matching our conceptual model of CD8⁺ T cell differentiation. These analyses illustrate how clonal data can be used to infer and validate differentiation trajectories *in vivo*.

We next compared isotype versus anti-PD-1 treatments and found no significant difference in overall clone-size distributions (Figure S6a; p=0.91). However, anti-PD-1-treated clones show fewer progenitors and a higher proportion of cycling cells (Figure S6b,c), suggesting that checkpoint blockade promotes differentiation. Clones with Jak3, Prdm1, and Trim28 gRNAs tend to have smaller clone sizes compared to NT controls (Figure S6d,e; p=0.0034 for Trim28). These observations corroborate our prior findings that Jak3 and Prdm1 function in T-cell activation and expansion. Ifngr1 showed relatively fewer anti-PD-1-treated clones than expected from the NT ratio (Figure S6f; p = 0.04).

Collectively, this barcoded Perturb-seq framework demonstrates a broadly applicable strategy for simultaneously perturbing and measuring the clonality of cells. We observe substantial heterogeneity amongst our clone sizes which allows us to determine clonal dynamics and infer cell type differentiation trajectories.

### Clonal counts are superior to cell counts in determining perturbation effects on T cell states

We next sought to determine whether cells harboring specific perturbations expanded or contracted (normalized to starting abundance) relative to controls. This is typically done by counting the number of guides, which corresponds to the number of cells with that guide. Importantly, we expect the number of cells with control guides should be proportional to the number of input cells with the same guides. However, one limitation is that clones arising from a single cell may expand differentially independent of guide identity. To test whether stochastic variation in clone expansion confounds the estimation of guide abundance in tumors after T cell entry and growth, we determined the relationship of initial guide abundance to either number of cells or number of clones containing each guide. If clones have a large spread in their cell expansion capabilities, then we expect a higher correlation of initial guide numbers to clone numbers rather than cell numbers. To evaluate this hypothesis, we calculated the correlation between initial guide abundance with the number of cells or clones recovered in the tumor.

Control guides showed weak correlation of cell numbers with their initial abundances (Spearman ρ=0.47, Pearson r=0.39) (Figure 4a). In contrast, when we related initial control guide abundance to the number of clones (instead of cells), we observed a much stronger correlation (Spearman ρ=0.86, Pearson r=0.94) (Figure 4b). We conclude that the number of clones per guide better reflects initial guide abundance (for guides with no expected biological effects).

**Figure 4:**
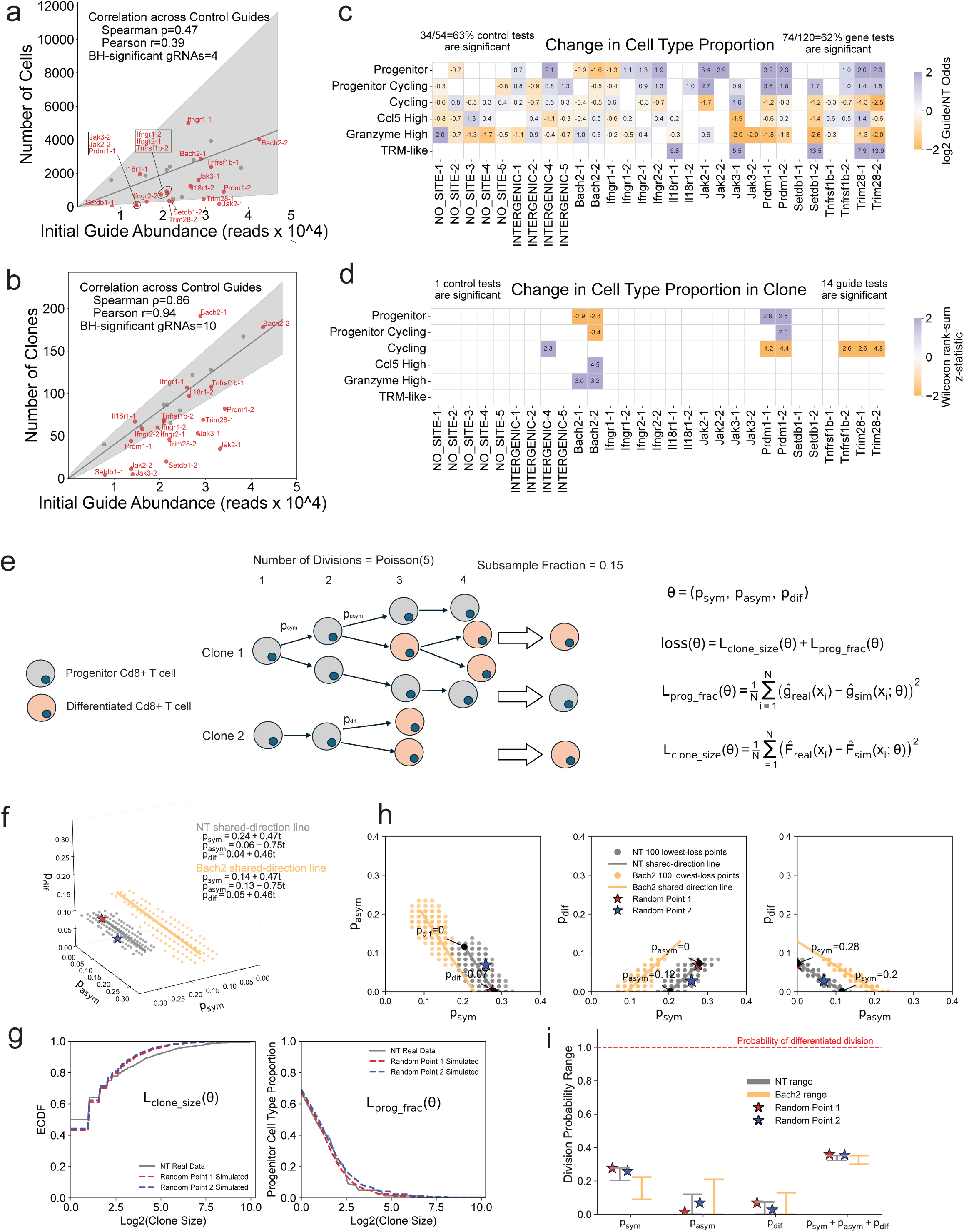
Clonal information improves ability to detect perturbation effects and discover rates of differentiation. (a) Scatter plot of the number of cells (y-axis) versus initial guide abundance (x-axis). Control guides are plotted in gray and gene-targeting guides in red. Gray line and shaded region represent the modeled relationship and 95% confidence interval relating initial guide abundance to counts, fit using control guides (Methods). Correlations across control guides only are reported on the plot. (b) Same as (a) but with the number of clones on the y-axis; control guides in gray, gene-targeting guides in red. (c) Heatmap showing enrichment or depletion of cell state frequencies resulting from gene perturbations. For each gRNA and each cell state, a Fisher’s exact test was applied to a two-by-two contingency table composed of the number of cells of a given state vs other states, for cells targeted with the gRNA in question vs control gRNAs. A gene-targeting gRNA was compared with all control gRNAs; a control gRNA was compared with all other control gRNAs excluding the gRNA in question. The plotted effect size is the log2 odds ratio (guide vs NT, with a +1 pseudocount), so positive values indicate higher odds of the cell state among guide-targeted cells when BH-adjusted (within cell type) FDR < 0.10. (d) Heatmap showing enrichment or depletion of cell state proportion in clones as a result of gene perturbations. For each gRNA and each cell state, we computed the frequency of cells within each clone that belong to the cell state in question, and used the Wilcoxon rank-sum test to compare these frequencies between the gRNA’s clones and control clones. A gene-targeting gRNA was compared with all control gRNAs; a control gRNA was compared with all other control gRNAs excluding the gRNA in question. Values are shown when BH-adjusted (within cell type) FDR < 0.10. (e) Schematic for modeling clonal data using simulations. In a simulation, each clone begins from a progenitor CD8+ T cell. First we assign the number of divisions based on a Poisson distribution with mean 5. At each division, the progenitor can self-renew to two progenitors with probability p_sym_, divide asymmetrically to one progenitor and one differentiated cell with probability p_asym_, or generate two differentiated cells with probability p_dif_; differentiated cells always divide into two differentiated cells. Lastly, for the 10-gene Perturb-seq analysis shown here, we subsample the expanded clone keeping each cell with probability 0.15. (f) The 100 lowest-loss parameter sets for NT (gray) and Bach2 (orange), obtained by grid search across (p_sym_, p_asym_, p_dif_), shown as a 3D scatter. Lines are fit through the NT and Bach2 lowest-loss points and constrained to share the same direction. Two NT parameter sets, randomly selected from the 100 lowest-loss points, are marked (red and blue stars). (g) Summary statistics used to compute loss between NT real data and simulations for the two parameter sets highlighted in (f). Left: empirical cumulative distribution function (ECDF) of log2(clone size) for NT real data (gray) and simulations under each parameter set (red and blue dashed). Right: fraction of progenitor-phenotype cells versus log2(clone size) for NT real and simulated clones, as in Figure 3d; curves fit by isotonic regression. (h) Orthogonal 2D projections (p_sym_ versus p_asym_; p_sym_ versus p_dif_; p_asym_ versus p_dif_) of the same lowest-loss points and shared-direction lines as in (f); the same two NT parameter sets highlighted in (f) are marked (red and blue stars). Endpoint labels indicate the values of the third parameter at the ends of each line, showing how it varies along the line. (i) Ranges for p_sym_, p_asym_, p_dif_, and p_sym_ + p_asym_ + p_dif_. Vertical bars denote permissible parameter values as determined by the shared-direction fit constrained by 0 ≤ p ≤ 1 and p_sym_ + p_asym_ + p_dif_ ≤ 1. Gray, NT; orange, Bach2. Stars mark the two NT parameter sets from (f). A horizontal red dashed line at 1.0 is shown as a reference representing that a differentiated cell always divides at each division cycle.

To identify guides with biological effects on cell abundance in the tumor, we tested for significant deviations of the number of cells or the number of clones per guide compared to control guides normalized to starting abundance. Using the cell-based approach, a large dispersion parameter (beta=0.4) was needed to account for the variability in control guide cell number, resulting in all perturbations falling within plausible bounds under the null hypothesis and thus deemed non-significant (Table S4.1), even for positive controls Bach2 and Prdm1. In contrast, using the clone-based approach, a significantly smaller dispersion parameter (beta=0.008) was needed to account for the variability in control guide clone number. Several gene perturbations fell outside the region of the null model confidence intervals and were statistically significant (Table S4.2), with Bach2 having more clones than expected, and Prdm1, Jak2, Setdb1, Trim28, and Jak3 having fewer clones than expected. Overall, by removing the effects of clonal stochasticity, clone counts are a better proxy for the effects of perturbations on T cell numbers than direct cell counts.

We next asked whether specific perturbations altered the frequency of particular cell states. Common approaches implicitly treat each cell as an observation of an independent perturbation event. However, all cells from a single clone arise from the same CRISPR edited cell.

Overconfidence and bias can arise when variable clone sizes, which can reflect stochastic differences in trafficking, growth, differentiation or death^10^, cause a small number of clones to dominate the cells assigned to a guide. In our data, large clones contained on average 42% of cells for a control guide despite there being 102 clones per control guide (Figure S7a), and could therefore dominate a perturbation and confound the observed perturbation effect.

Consistent with this, the Fisher’s test which assumes that cells are independent observations identified numerous significant differences even between control gRNAs: 34 of 54 control-versus-control tests (63%) were called significant at BH-adjusted FDR < 0.10 (Figure 4c; Figure S7b). Because all of these should be null comparisons, this represents an unacceptably high false-positive rate. Moreover, this fraction was nearly identical to the rate observed for gene-targeting gRNAs (74 of 120 tests; 62%), leaving little confidence that the apparent perturbation effects exceed the background false-positive rate (Figure 4c).

To minimize these issues, we next looked at the shifts of phenotypic distributions at the clonal rather than cellular level. Using the clonal information, we developed an analogous test that compares clones of a perturbation against control clones (or clones from one control against clones from all other controls). This clonal approach treats each clone as an independent observation, more closely reflecting the data’s independence structure—each clone originates from a single edited cell whose descendants arise through subsequent divisions. We found that this clonal test greatly reduced false positives among control guides, leaving only 1 of 54 control-versus-control tests significant at BH-adjusted FDR < 0.10, while still recovering the expected increases in progenitor frequency for Bach2 and Prdm1 positive controls (Figure 4d; Figure S7c). This improved specificity suggests the clonal test is more accurate and reliable than the cell-based test. Using the clonal test, we also observed that Trim28 gRNA resulted in decreased frequency of cycling cells (Figure 4d; Figure S7d). A caveat of the clonal analysis is a reduced number of observations, which can reduce sensitivity; for example, the clonal analysis did not identify the TRM-like cluster or the Progenitor cluster as significantly associated with Trim28 gRNA—likely due to few clones in that cluster—while the cell-based test did (Figure 4c,d).

As a quality-control step for analysis presented in Figure 4a–d, we excluded one intergenic control gRNA that behaved as an outlier (and overlapped a chromatin-accessible (ATAC-seq) peak in our multiomic experiment, see Figure 6 below); however, analyses including all control guides yield similar conclusions (Figure S8a-d).

### Clonal analysis enables inference of cell division rates and patterns

In our clonal dataset, each clone represents a single snapshot of clonal progression reflecting the history of expansion, differentiation, and death of cells arising from a single barcoded naive T cell. However, clones are not synchronized and thus each snapshot captures clones at different times after initial priming and entry into tumors. Prior tumor studies showed that progenitor exhausted CD8 TILs persist and differentiate into terminally exhausted TILs.^2^ We reasoned that the collection of clonal snapshots would encode information about the underlying rates of progenitor self-renewal and differentiation. To test this, we constructed a stochastic simulation of progenitor division and differentiation and identified parameter sets for which simulated clones recapitulate the empirical relationships between clone size and cell-state composition.

Our simulation of cell division tracks only two cell states: progenitor cells (Progenitor and Progenitor Cycling) and differentiated cells (Methods). Each simulated clone begins from a single progenitor, and at each cycle of division, a progenitor can undergo: (1) symmetric self-renewal (with probability p_sym_) to produce 2 progenitors; (2) asymmetric division (with probability p_asym_) to produce 1 progenitor and 1 differentiated cell; (3) symmetric differentiation (with probability p_dif_) to produce 2 differentiated cells; (4) no division (with the remaining probability). We assume that differentiated cells divide to produce 2 differentiated cells in every cycle. For the 10-gene Perturb-seq analysis shown in Figure 4, after clonal expansion each cell is retained independently with probability 0.15 to mimic incomplete capture (Figure 4e). The number of division cycles is drawn from a Poisson distribution with mean 5 to match the clone size distribution.

For any choice of parameters (p_sym_, p_asym_, p_dif_), this model yields simulated clones with a distribution of cell numbers and states. To find parameters that best match the real data, we compared simulated and observed clones using a loss function representing the deviation of two summary statistics: (i) the distribution of clone sizes and (ii) the fraction of progenitor cells as a function of clone size (Figure 4g). We found that a range of parameters could fit the data well (Figure 4f,g). When applied across different gRNAs, the curve fits had lowest loss for NT and Bach2 (Figure S9a,b), likely reflecting their larger sample size (number of clones); thus, we were most confident in these simulations and restricted further analysis to these perturbations.

To characterize the structure of parameter sets consistent with the data, we evaluated the loss across a dense three-dimensional grid of (p_sym_, p_asym_, p_dif_) values and visualized the 100 lowest-loss points. Both NT and Bach2 points formed linear trends that ran in the same direction through the (p_sym_, p_asym_, p_dif_) space but were offset from one another (Figure 4f,h). Along this direction, the parameters change in a coordinated manner: for each unit of movement, p_sym_ increases by +0.47, p_asym_ decreases by –0.75, and p_dif_ increases by +0.46. This pattern reflects the trade-offs imposed by the model—higher symmetric self-renewal must be offset by lower asymmetric division and greater symmetric differentiation to maintain the observed progenitor fraction and clone-size distribution. These compensations yield many parameter triplets with essentially indistinguishable loss (Figure 4h).

For both NT and Bach2, we found that the rate of symmetric self-renewal was nonzero and much higher than the rate for asymmetric division and symmetric differentiation (Figure 4i) which explains how a cell maintains a population of progenitors. Additionally, the aggregate probability that a progenitor divides in a given cycle (p_sym_ + p_asym_ + p_dif_) is on the order of 0.3–0.4 (Figure 4i), well below 1, which is consistent with the fact that differentiated cells, unlike progenitors, divide into two differentiated cells every division cycle. Indeed, differentiated expansion outpaces progenitor renewal resulting in the clonal expansion of differentiated cells.

Comparing perturbations, Bach2’s lowest-loss solutions are shifted relative to NT toward lower p_sym_ and higher p_asym_ / p_dif_ (Figure 4h,i), consistent with Bach2 loss promoting differentiation because Bach2 normally enforces stem-like CD8 T cell programs during chronic infection and cancer.^23^

Together, these analyses show how clonal data, when coupled with a cell-division simulation and a curve-matching loss, allows estimation of plausible ranges for natural division rates and how these change across perturbations.

### Trim28 and Setdb1 deficiency restricts T cell differentiation and leads to a TRM-like phenotype

In our *in vivo* CRISPR screens (Figure 1h), we observed that Trim28 and Setdb1 gRNA resulted in higher numbers of Tcf7 expressing cells. We further confirmed this with our Perturb-seq experiment finding that Trim28 and Setdb1 gRNA resulted in fewer differentiated cells and more progenitor cells and a reduction in cell numbers. Strikingly, we also observed that cells with Trim28 or Setdb1 gRNAs selectively formed a TRM-like phenotype. Furthermore, both genes are known to form a shared epigenetic complex.^26^ Because the initial experiment was underpowered for clonal testing, we performed the higher-resolution, clonally barcoded Trim28/Setdb1 Perturb-seq experiment focused on Trim28 and Setdb1 perturbations. We pooled ten gRNAs against Trim28, ten against Setdb1, and six NO-SITE controls; Trim28-1/Trim28-2 and Setdb1-1/Setdb1-2 are the same guide sequences used in the 10-gene Perturb-seq screen. We transduced naive Cas9 x OT1 CD8+ T cells, and adoptively transferred them into B16-OVA–bearing mice. Mice then received either isotype or anti-PD-1 therapy. We subsequently recovered donor OT1 CD8+ T cells from tumor and draining lymph nodes and profiled each cell’s transcriptome together with its clonal barcode, guide assignment, mouse hashtag, and location from different sequencing lanes (tumor or lymph node) (Figure 5a).

**Figure 5:**
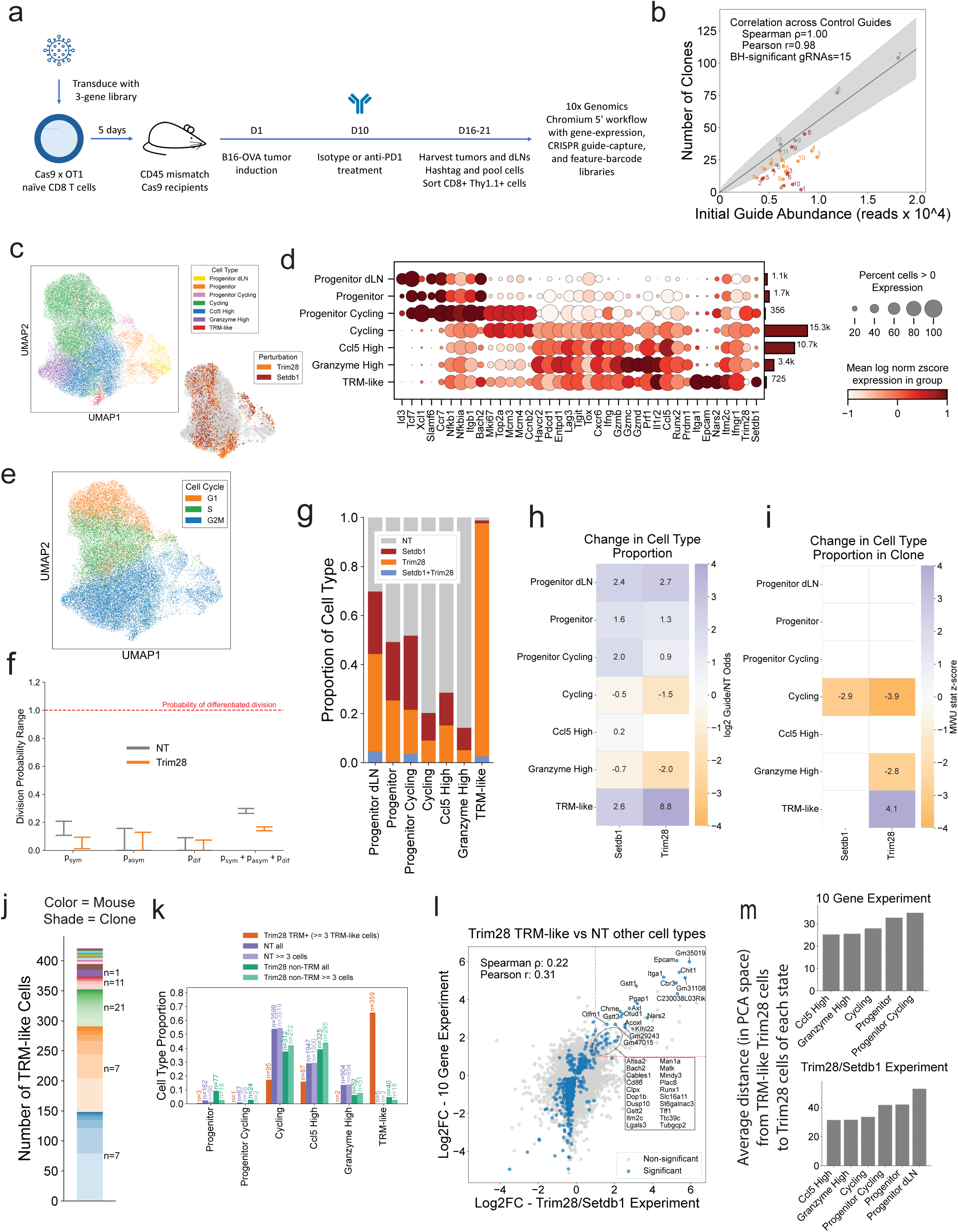
Trim28 and Setdb1 perturbations restrict T cell differentiation within tumors and bias toward a novel TRM-like trans criptional state. (a) Schematic of the experimental design for clonally barcoded Perturb-seq approach targeting Trim28 and Setdb1. Naive Cas9 x OT1 CD8+ T cells were transduced with a construct library containing 10 Trim28 guides, 10 Setdb1 guides, and 6 NO-SITE control guides, then adoptively transferred into tumor-bearing mice. Recovered donor cells from tumor and lymph node were profiled for their transcriptome, mouse hashtag assignments, CRISPR perturbations, and clonal barcodes via single-cell approaches. (b) Scatter plot relating recovered clone counts (y-axis) to initial guide abundance (x-axis); control guides are plotted in gray and gene-targeting guides are colored by perturbation. Labels indicate guide number. Gray line and shaded band show the negative-binomial fit and 95% predictive interval learned from NT controls (analogous to Figure 4b). (c) UMAP representation illustrating cell states based on annotation of clusters obtained by Leiden clustering of the kNN graph of cell-cell transcriptomic similarities (left), and highlighting Trim28 versus Setdb1 perturbations (right). (d) Dot plot visualizing the expression levels of key markers across identified cell states across all cells. Dot size represents the percentage of cells expressing each marker, while color intensity shows z-score normalized log expression levels. The bar plot on the right of the dot plot shows the number of cells per cell type. (e) UMAP plot displaying cell cycle phases. (f) Inferred parameter ranges (analogous to Figure 4i) for p_sym_, p_asym_, p_dif_, and their sum using the clonal differentiation rate simulations. NT is shown in gray and Trim28 in orange. (g) Stacked bar chart showing the relative abundance of perturbations within each cell state. (h) Cell-based heatmap showing change in cell type proportion per perturbation, comparing each perturbation to NT cells with 2×2 Fisher’s exact tests (analogous to Figure 4c). (i) Clone-based heatmap showing change in cell type proportion per perturbation, comparing each perturbation’s clones to NT clones using Wilcoxon rank-sum tests (analogous to Figure 4d). (j) Stacked bar plot showing the number of TRM-like cells per clone, ordered by mouse. Each color denotes a different mouse and progressively darker shades within a color correspond to distinct clones from that mouse. n labels (shown for the top five mice) indicate the number of TRM-like clones contributing for that mouse. (k) Cell type proportion of cells from different sets of clones; Trim28 TRM+ clones are those with at least 3 TRM-like cells; Trim28 non-TRM clones are Trim28 clones with fewer than 3 TRM-like cells. Comparison groups include all NT cells, NT cells from clones with ≥3 cells, cells from Trim28 non-TRM clones, and cells from Trim28 non-TRM clones with at least 3 cells. (l) Scatter plot comparing log2-fold changes (log2FC) for TRM-like Trim28 gRNA cells versus non–TRM-like NT cells across the 10-gene Perturb-seq (x-axis) and Trim28/Setdb1 Perturb-seq (y-axis) experiments. Highlighted genes are the TRM-like signature genes, defined as genes with Benjamini-Hochberg adjusted P < 0.05 in both experiments and log2FC > 1 in both experiments (dotted red lines). (m) Bar graph showing, for each other cell state, the mean Euclidean distance in PCA space (first 50 PCs) computed across all TRM-like Trim28 cells paired to all Trim28 cells of that state within each experiment.

Consistent with our earlier analyses (Figure 4b), clone-based analysis more accurately captured the effect of perturbations on cell number (Figure S10a). Clones with Trim28 or Setdb1 gRNAs were recovered less frequently than expected relative to their initial guide abundance (Figure 5b). Setdb1 guides showed a more negative average NB standardized residual (−3.36) than Trim28 guides (−2.63), indicating stronger depletion of recovered cell number relative to the control-fit null model. We also found that Setdb1- and Trim28-targeted clones were significantly smaller than NT controls, with Setdb1 producing the larger reduction in clone size compared to Trim28 (Figure S10b). This suggests that Trim28 and Setdb1 perturbation constrains clonal recovery and expansion.

After clustering the single-cell data from donor cells, we found that donor cells from the tumor and lymph node separated into different clusters. The lymph node donor cells expressed stem-like T cell genes, and thus we termed them Progenitor dLN. The tumor donor cells were clustered in the same major T-cell states as before—Progenitor, Progenitor Cycling, Cycling, Ccl5 High, Granzyme High, and TRM-like (Figure 5c-e; Figure S11a). We did not observe any clones that were shared between dLN and tumor, possibly due to low guide recovery – just 10% of LN (28% for tumor) cells had gRNAs/clonal barcodes – and to undersampling of cells. Clones from dLN had sizes of just one or two cells.

We applied our clonal modeling approach from Figure 4e-i to NT and Trim28 clones (Figure 5f). As before, the model was able to recapitulate key features of clonal behavior, including both the distribution of clone sizes and the progenitor composition within clones (Figure S12a). We found not as strong a fit for Setdb1 clones, possibly due to fewer clones (Figure S12a). For NT and Trim28, we found a linear manifold of well fitting parameter sets where higher symmetric self-renewal must be offset by lower asymmetric division and greater symmetric differentiation (Figure S12b,c). Compared to NT, our simulations predict that Trim28 has a much lower rate of symmetric self-renewal while the other rates of division stay relatively constant (Figure 5f).

We next evaluated whether Trim28 and Setdb1 depletion affected cell state frequencies (Figure 5h,i). Using a Fisher’s exact test for changes in cell type proportion, we found that cells with Trim28 and Setdb1 gRNA had more Progenitors (in dLN and tumor) and fewer Cycling and Granzyme High states (Figure 4c; Figure 5h). A Wilcoxon rank-sum–based analysis testing for differences in cell type proportion within clones (similar to Figure 4d) showed significantly fewer Cycling cells per clone upon loss of Trim28 or Setdb1 (Figure 5i), implying that both genes are necessary for proliferative expansion. We also found that Trim28 clones tended to have fewer Granzyme High cell states and were enriched for the TRM-like state (Figure 5i).

The TRM-like state was formed by cells with Trim28 gRNAs or NG cells, but not cells with NT control guides (Figure 5g; Figure S11b). Thus, we infer that the NG cells in the TRM-like state likely had the Trim28 gRNA but we were unable to detect gRNAs possibly due to low guide recovery. We found that multiple Trim28 clones (Figure 5j) and guides (Figure S11c) contributed to the TRM-like cells providing reproducible evidence that the Trim28 gRNA leads to TRM-like state.

Amongst expanded Trim28 gRNA clones, we observed some clones having a high proportion of cells in the TRM-like state and other clones without the TRM-like state (Figure S11d). We suspect one reason for this is because CRISPR may introduce functionally neutral edits or may fail to edit. Thus, we sought a more stringent approach to confidently identify clones with deleterious edits in Trim28. In parallel, we asked which cell states are preferential within Trim28 clones with a high TRM-like population. Since only TRM-like cells within Trim28 clones exhibited strong perturbation signatures, we defined Trim28 TRM+ clones as those containing at least 3 TRM-like cells. We compared these against NT and Trim28 non-TRM clones using both all clones and a ≥3-cell cutoff for those groups (Figure 5k). Within the cells of the Trim28 TRM+ clones, most cells were found in the Cycling or Ccl5 High states, while Granzyme High cells were notably absent (Figure 5k). This pattern suggests that Trim28 deficiency and TRM-like formation allows progression to intermediate effector states but restricts full differentiation into the terminal Granzyme High state. Using the clone-size-normalized cell type enrichment analysis as in Figure 3f on Trim28 clones, we found that TRM-like state may be at some intermediate differentiation level between Ccl5 High/Cycling and Granzyme High (Figure S11e).

Next, we aimed to better characterize the TRM-like state. We calculated the differential expression of Trim28 TRM-like cells against all other T-cell states in both the 10-gene Perturb-seq and Trim28/Setdb1 Perturb-seq datasets (Figure 5l). We found a strong global correlation of logFC gene expression changes between datasets. We define a TRM-like signature as genes with Benjamini-Hochberg adjusted P < 0.05 and log2FC > 1 in both experiments (Table S5.2).

Itga1 again featured prominently, reflecting a potential role in tissue residency. We also found Epcam upregulated in TRM-like states across experiments. Although Epcam is classically characterized as an epithelial cell adhesion molecule, its upregulation in TRM-like cells may indicate an adhesion or retention program that promotes CD8⁺ T cell persistence within tissues. We found that TRM-like cells were more similar to the differentiated cell states than to the progenitor populations (Figure 5m).

In this experiment and in contrast to our first experiment, cells with Setdb1 gRNAs did not contribute to a discrete TRM-like state (Figure 5g). Despite this, all 9 Setdb1 guides with more than 10 cells in our second experiment showed significantly higher expression of genes that were differentially expressed between TRM-like and NT cells (Figure S11f). Thus, we infer that some Setdb1 cells were intermediate between the Trim28 TRM-like state and NT cells. We also found correlation of Setdb1 vs NT logFC and Trim28 vs NT logFC for both experiments (Figure S11g). This suggests that Trim28 and Setdb1 regulate overlapping transcriptional programs associated with a TRM-like phenotype.

Overall, these data suggest that Trim28 and Setdb1 play overlapping roles in T-cell fate decisions within the tumor microenvironment.

### Trim28 loss increases chromatin accessibility at TRM-like gene loci such as Itga1

Informed by the Trim28-dependent phenotypes observed in our previous experiments (Figure 5) and the fact that Trim28 is known to influence chromatin accessibility through forming a complex with Setdb1 (a chromatin repressor)^26,27^, we next performed a multiomic single-cell RNA-seq + ATAC-seq analysis to explore how Trim28 regulates chromatin accessibility in donor CD8+ T cells. We profiled congenically distinguishable donor Trim28-1 gRNA and NT control OT-I cells recovered from the same mixed competitive-transfer B16-OVA tumor experiment approximately two weeks after tumor implantation. We then clustered cells using scATAC-seq signals (Figure 6a). In our initial clustering attempt we found significant batch effects across conditions, so we performed integration with Harmony.^28^ We annotated the clusters based on the scRNA-seq expression of marker genes where we found the same cell types as before (Figure 6a; Figure S13a,b; Table S6.1). TRM-like cells in this experiment scored highly based on the TRM-like signature from our previous experiments (Figure 5l). This analysis demonstrates that TRM-like cells can be distinguished in a data-driven manner in not just RNA space but also chromatin-accessibility (ATAC) space even after integration. In the TRM-like state we identified 40 donor cells with Trim28 gRNA and 2 donor cells from control, providing further evidence that Trim28 gRNA drives cells to this state. In both scRNA-seq and scATAC-seq PCA spaces, TRM-like cells were closer to differentiated states than to the Progenitor state, with Progenitor showing the largest mean distance in both modalities (Figure S13c), similar to our earlier Perturb-seq result (Figure 5m). Trim28 cells were also enriched for the Progenitor state relative to controls in the multiome scRNA-seq dataset (Figure S13d), similar to the Perturb-seq result (Figure 4c; Figure 5h).

**Figure 6.**
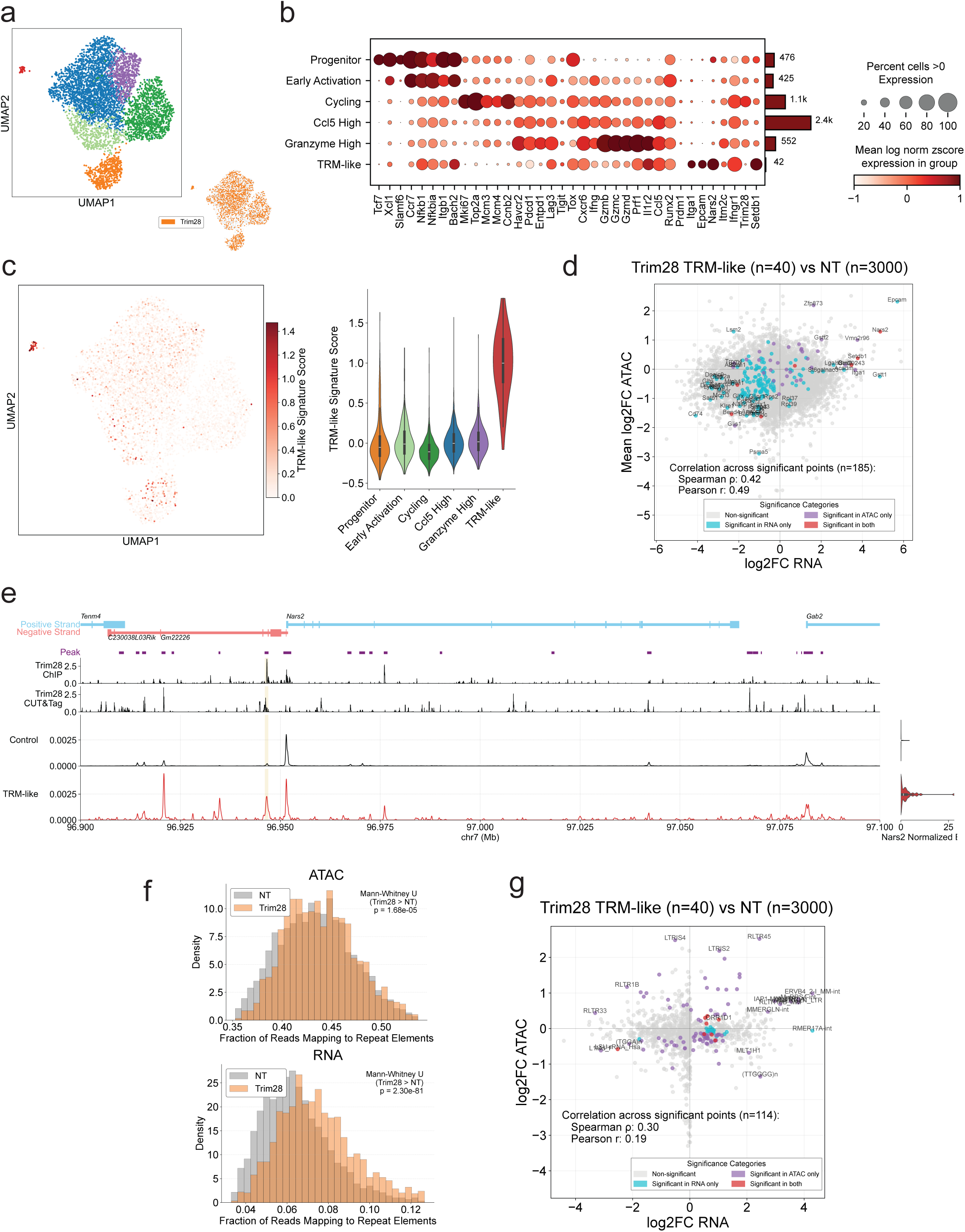
Multiomic single-cell RNA-seq + ATAC-seq analysis of Trim28-deficient cells reveals a TRM-like accessibility signature and changes in repeat element accessibility and gene expression. (a) Harmony-corrected UMAP visualization derived from single-cell ATAC-seq profiles of donor CD8+ T cells from control or Trim28 gRNA cells. Donor cells were isolated from tumors, and clustering was performed based on ATAC-seq signals. In the bottom right, the UMAP is colored by cells with Trim28 gRNA. (b) Dot plot summarizing scRNA expression for the same cells profiled in parallel, highlighting key markers that distinguish each cell state. Dot size represents the percentage of cells expressing each marker, while color intensity shows z-score normalized log expression levels. The bar plot on the right of the dot plot shows the number of cells per cell type. (c) TRM-like gene signature scores (as defined in Figure 5l) used to color the ATAC-seq-based UMAP (left) and shown as violin plots (right). (d) Scatter plot relating changes in accessibility and gene expression comparing Trim28 TRM-like cells to non-TRM-like control cells. Each point represents a gene; ATAC-seq log2FC is computed by averaging log2FC across peaks assigned to that gene by nearest-gene mapping. Genes were classified as ATAC-significant when at least one assigned peak had BH-adjusted P < 0.05. (e) Genome browser tracks showing Trim28 ChIP-seq and CUT&Tag binding and accessibility around the Nars2 locus in Control and TRM-like cells. The shaded region marks the Nars2-assigned ATAC-seq peak that was significant in the Figure 6d comparison. The right panel shows the corresponding normalized RNA-seq expression of Nars2. (f) Histograms quantifying the fraction of reads mapping to repeat elements for Trim28 gRNA and control cells in both ATAC-seq (top) and RNA-seq (bottom) modalities. (g) Scatter plot relating changes in accessibility and gene expression for repeat eleme nts comparing Trim28 TRM-like cells to non-TRM-like control cells.

The accessibility and expression changes in Trim28 gRNA cells correlated well overall (Figure 6d; Table S6.2). Trim28-deficient TRM-like cells had an increase in expression of *Nars2*, *Itga1* (Figure S14a) and *Setdb1* (Figure S14b) and an increase in accessibility at these loci. At the *Nars2* locus (Figure 6e), a nearby peak had significantly higher accessibility in TRM-like cells together with higher RNA expression (Figure 6e; Table S6.2). Consistent with a potential direct relationship, the highlighted accessible region overlaps reported Trim28 binding tracks.^27^

Given Trim28’s known interaction with Setdb1 (a histone methyltransferase typically associated with repressive H3K9me3 marks) and the known role of this pathway in silencing repetitive elements^26,29,30^, we next examined accessibility and expression of repeat regions. Strikingly, Trim28 gRNA cells displayed significantly higher accessibility and transcription at these repeat loci (Figure 6f), suggesting that Trim28 normally restricts aberrant chromatin opening and RNA production from repeats. Repeat elements that were more accessible tended to show higher RNA levels (Figure 6g; Table S6.3).

Collectively, these multiomic data highlight accessibility profiling of the TRM-like cells. Furthermore, Trim28 gRNA affects accessibility of TRM-like genes and repeat elements in a tumor-infiltrating CD8+ T cell population.

### Trim28 deficiency increases proportion of Tcf7+ T cells, alters tissue distribution, and increases relative tumor recovery with anti-PD-1 therapy

To better understand the impact of Trim28 deficiency on the T cell state distribution in tumors, and to specifically determine the fraction of Trim28 knockout TILs in a Tcf7-positive state, naive CD8 T cells were transduced with either Trim28 or NT guides and transferred into recipient mice implanted with B16-OVA tumors. When tumors were harvested just prior to euthanasia endpoint, flow cytometry showed that a higher percentage of Trim28 knockout TILs fell within the intracellular TCF7+ gate compared with matched NT TILs (Figure 7a). This validation result was consistent with the Tcf7 CRISPR screen and *in vivo* Tcf7 sort-screen results that nominated Trim28 for follow-up (Figure 1h), and with Perturb-seq analyses showing that Trim28 perturbation shifted donor T cells toward higher progenitor proportion (Figure 4c,d; Figure 5g-i).

**Figure 7:**
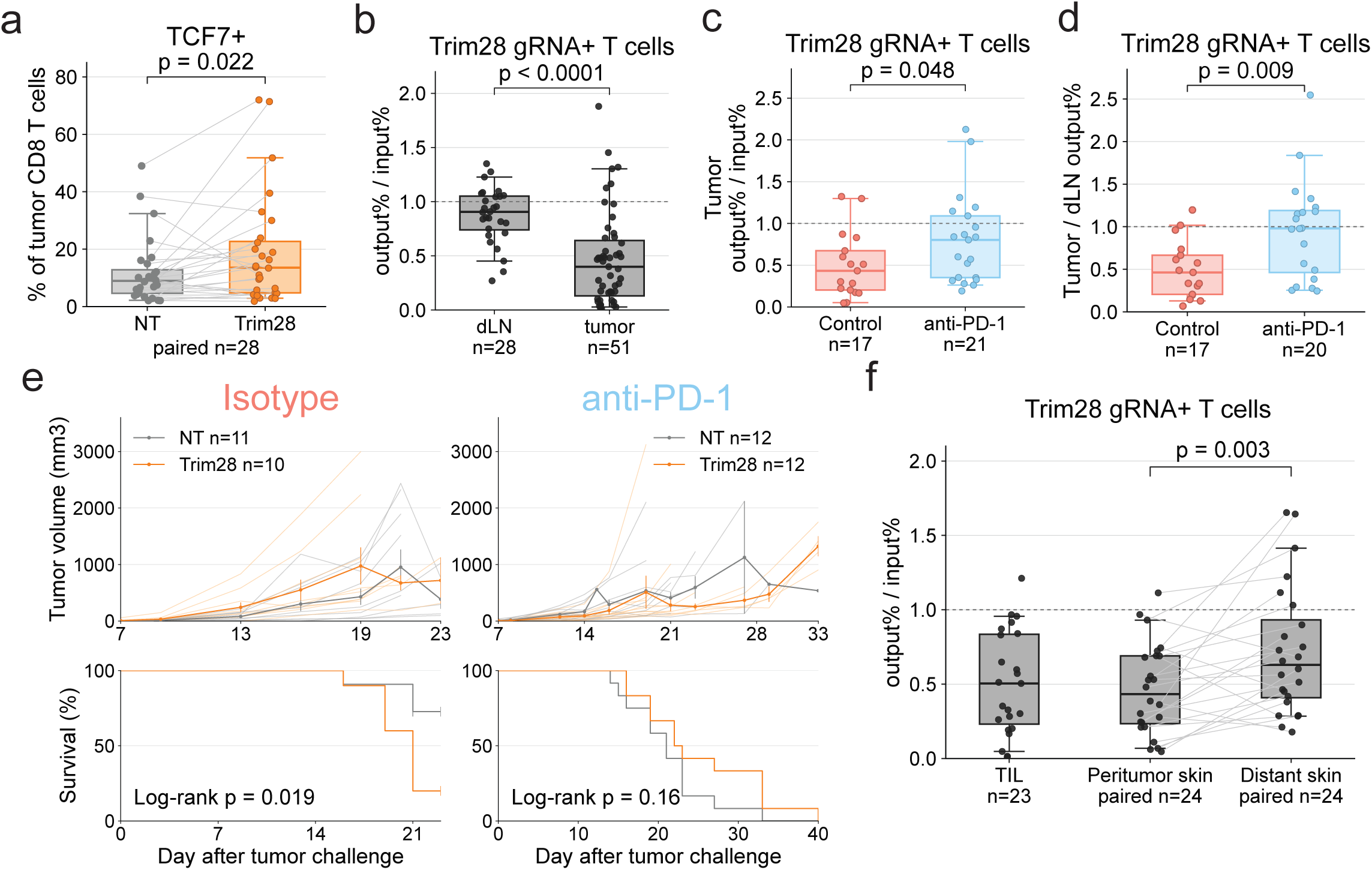
Functional effects of perturbing Trim28 *in vivo*. (a) Flow cytometry quantification of the percentage of NT or Trim28 gRNA donor TILs gated as intracellular TCF7+ after competitive transfer of non-targeting (NT) and Trim28 gRNA cells. n is the number of mice. The P value shown was calculated with a two-sided Wilcoxon signed-rank test. (b) For draining lymph nodes (dLNs) and tumors after competitive transfer of NT and Trim28 cells, flow cytometry quantification of the percentage of recovered (output) donor cells that were Trim28 deficient divided by the percentage of cells prior to transfer (input) that were Trim28 deficient. n is the number of mice. The P value shown was calculated with a two-sided Mann-Whitney U test comparing dLN and tumor values as unpaired groups. (c) Percentage of recovered (output) donor cells in the tumor that were Trim28 deficient divided by the percentage of cells prior to transfer (input) that were Trim28 deficient after isotype control or anti-PD-1 treatment. Values were measured by flow cytometry in congenic-marker experiments and by guide sequencing in guide-count experiments. n is the number of mice. The P value shown was calculated with a two-sided Mann-Whitney U test. (d) Percentage of recovered (output) donor cells in the tumor that were Trim28 deficient divided by the percentage of recovered (output) donor cells in the dLN that were Trim28 deficient after isotype control or anti-PD-1 treatment. Values were measured by flow cytometry in congenic-marker experiments and by guide sequencing in guide-count experiments. n is the number of mice. The P value shown was calculated with a two-sided Mann-Whitney U test. (e) Tumor growth and endpoint-free survival in mice receiving OT1 T cells transduced with NT or Trim28 gRNAs, with isotype control or anti-PD-1 treatment. In tumor-growth plots, light lines show individual mice, dark lines show group means, and vertical bars show s.e.m. at each time point. n is the number of mice. The endpoint-free survival P values shown were calculated with two-sided log-rank tests. (f) Percentage of recovered (output) donor cells from the tumor or areas surrounding the tumor that were Trim28 deficient divided by the percentage of cells prior to transfer (input) that were Trim28 deficient after competitive transfer, quantified by guide sequencing and restricted to mice with matched peritumor and distant skin samples. Paired n is the number of matched peritumor-distant skin sample. The P value shown for the matched peritumor-distant comparison was calculated with a two-sided paired sign-flip test.

We next performed competitive transfer experiments to assess abundance of Trim28-deficient vs NT cells in draining lymph nodes (dLNs) and tumors. We quantified cell numbers by flow cytometry using congenic markers to track proportion of Trim28 KO vs NT T cells prior to transfer (starting ratio) and when tumors were harvested. We did not observe a significant difference in Trim28 proportion (i.e. of the mix of Trim28 and NT cells) in tumor draining LNs compared to the starting ratio. In contrast, we observed a decrease in the proportion of Trim28 cells in the tumor, consistent with the lower abundance of Trim28 KO cells relative to NT cells observed in the single-cell screens *in vivo* (Figure 4b). We then treated mice with anti-PD-1 or isotype control using the same competitive transfer approach, combining flow cytometry-based congenic-marker experiments with guide-sequencing measurements. We found an increase in the proportion of Trim28 KO cells for anti-PD-1 treatment compared to control in the tumor (Figure 7c). As before (Figure 7b), we observed a decrease in the proportion of Trim28 cells in the tumor relative to the dLN in isotype treatment (Figure 7d). After anti-PD-1 treatment, the relative proportions of Trim28 KO cells in the tumor and dLN increased (Figure 7d). Additionally within the anti-PD-1 group, the fraction of recovered Trim28-deficient cells in tumors was not significantly different from the fraction in matched dLNs (P = 0.62, two-sided Wilcoxon signed-rank test versus 1). These data suggest that anti-PD-1 treatment increases the relative recovery of Trim28-deficient cells in tumors.

In an effort to quantify the potential role of Trim28 KO cells in mediating tumor cell death, mice were injected with B16-OVA tumors and OT1 T cells transduced with either NT or Trim28 sgRNAs (Figure 7e). No tumor volume control or survival benefits were observed in the Trim28 KO group as compared to the NT group, and ICB did not reverse this trend.

Based on the TRM-like state observed with Trim28 perturbation, we tested whether Trim28 loss altered tissue distribution in a manner consistent with changes in trafficking, retention, or residency (Figure 7f). In a competitive transfer experiment, naive CD8 OT1 T cells transduced with Trim28 and NT sgRNAs were mixed and introduced *in vivo* by tail vein injection. B16-OVA cells were injected subcutaneously and after tumor growth for ∼2 weeks, skin samples were harvested underlying, adjacent to, and on the opposite flank from the tumor site. gRNAs were recovered from the skin samples by tissue digestion and PCR, and the ratio of Trim28 to NT guide normalized to the starting ratio was measured by guide sequencing. We observed a higher proportion of Trim28 KO cells in the skin farther from the tumor implant site, supporting the potential role of Trim28 in T cell tissue distribution and residence (Figure 7f).

## Discussion

We developed a naive-state-preserving CRISPR screening platform to identify regulators of CD8 T cell differentiation in tumors, enabling us to find genes controlling Tcf7 expression. By combining *in vitro* genome-wide screens, focused *in vivo* screens, and clonally barcoded Perturb-seq, we were able to measure how perturbations shape T cell abundance, clonal expansion, and differentiation state. This framework nominated Trim28 as a regulator that restrains TCF7-positive and TRM-like states while supporting the expansion and terminal differentiation of tumor-infiltrating CD8 T cells. Together, these results identify Trim28 as a regulator of CD8 T cell fate and show that clonal information can improve interpretation of single-cell CRISPR screens *in vivo*.

The integration of clonal barcoding into our Perturb-seq framework enabled analyses of T cell responses to tumors that were difficult to perform reliably with standard single-cell genomics approaches. In clonally resolved analyses, we observed substantial stochasticity both in clonal expansion and functional fates, even among clones with control guide RNAs. This clonal resolution captured an important layer of biological variance that is typically missing, averaged out or ignored as noise in standard scRNA-seq, scATAC-seq, multi-omic or Perturb-seq datasets.

Critically, clonally resolved statistical analysis provided a more rigorous foundation for assessing perturbation effects. In our Perturb-seq experiments, each guide-barcode integration event marks a single naive cell that later expands into many descendants before endpoint analysis; therefore, the independent unit of effect is the clone, not the cell. Thus in cell-based analysis, control guides can look similar to perturbation guides or could be confounded by the effect of “jackpot” clones, large stochastically expanded populations that typically dominate single-cell profiling and could generate false positives in cell-level analysis. In clone-level analysis, we mitigated these confounding effects. However, cell-based tests have higher apparent sample size and thus greater sensitivity, but violate independence; clone-based tests respect independence at the cost of a reduced number of observations.

Our approach allowed us to decouple perturbation effects on clonal persistence (number of clones that survive), proliferative capacity (clone size distribution) and functional fate (within-clone cell state composition) in tumor-targeting T cells, which are axes of clonal response ambiguated by cell-level analysis. In unperturbed clones we observe that increasing clonal expansion as measured by clone size and cell number is correlated with more differentiated cells. We find that perturbations did not affect just clonal expansion or differentiation alone. Prdm1, Jak2, Trim28, Setdb1, and Jak3 resulted in fewer cell numbers, indicating reduced clone size and reduced differentiation whereas Bach2 resulted in larger clones and more differentiation; these findings were not significant with cell based statistical analyses.

Clonal information also allowed us to infer plausible ranges of division and differentiation parameters under a simplified branching model. Although these estimates are model-dependent rather than direct measurements, the best-fitting parameter ranges suggested that symmetric progenitor self-renewal was consistently nonzero and greater than either asymmetric division or symmetric differentiation, consistent with maintenance of a progenitor pool during the tumor response. At the same time, the aggregate probability of progenitor division was lower than the model-imposed division probability for differentiated cells, providing a quantitative explanation for why larger clones were dominated by differentiated states. Comparing across perturbations, Bach2 loss was consistent with increased differentiation, whereas Trim28 loss was consistent with reduced progenitor self-renewal and constrained effector accumulation.

Our results have broader implications for the single-cell genomics field. Most standard scRNA-seq computational analysis pipelines implicitly assume that individual transcriptomic profiles are independent observations. However, our data reveal a high degree of transcriptomic interdependence among cells derived from the same recently expanded clone, creating bias that can distort definitions of phenotypes and trajectories and affect differential gene expression analysis and marker gene detection. This dependence is particularly relevant in systems defined by continuous cell expansion and differentiation such as development, hematopoiesis, immune response and tumorigenesis. As single-cell technologies continue to evolve, our results underscore the necessity of moving toward clonally or lineage aware experimental and computational approaches. Further research is needed to revisit the standard cell independence assumptions and develop better recommendations and standard practices for single-cell genomic analysis.

Across the *in vitro* and *in vivo* screens, we repeatedly recovered established regulators of Tcf7 and CD8 T cell differentiation, including Bach2 and Prdm1, validating the screening strategy.^23–25^ The screens also nominated IFNγR signaling, which was recently shown to regulate intratumoral stem-like CD8 T-cell maintenance and clonal diversity.^31^ In tumors, naive tumor-specific CD8 T cells undergo rapid clonal expansion and differentiate along an exhaustion hierarchy from Tcf7+ progenitor-exhausted (stem-like) cells toward Tcf7- terminally exhausted states.^2^

Trim28 scored modestly *in vitro* but showed stronger phenotypes *in vivo*, where Trim28 perturbation biased cells toward TCF7-positive progenitor states, reduced recovered cell numbers and clone sizes, and enriched for a TRM-like state. This TRM-like state expressed adhesion and residency-associated genes, including Itga1 and Epcam, and appeared more differentiated than the progenitor states, suggesting that Trim28 loss redirects some clones toward an alternative differentiated state rather than simply preserving an undifferentiated progenitor pool. TRM-like programs are clinically relevant because tissue-resident memory phenotypes have been linked to stronger cytotoxic responses or improved prognosis in multiple human tumor settings.^32–34^ The enrichment of Trim28-deficient cells in skin distant from tumor implant site is consistent with altered trafficking, retention, or survival in non-tumor tissue, but direct residency assays will be needed before concluding that Trim28 loss establishes bona fide tissue residence. Multiome profiling shows increased accessibility at TRM-like loci (e.g., *Itga1*, *Nars2*) with concordant RNA upregulation in Trim28-deficient cells, and broad de-repression of repeat elements at both ATAC-seq and RNA-seq levels. These differentiation and chromatin changes align with prior work linking Trim28 and Setdb1 to T-cell activation, differentiation, CD8 T-cell activation-associated chromatin remodeling, and repeat-element repression.^27,35–38^ Together with those studies, our findings extend the Trim28/Setdb1 chromatin regulatory axis to tumor-infiltrating CD8 T cells, where Trim28 perturbation couples altered chromatin and repeat-element regulation with enrichment of TCF7-positive and TRM-like states and reduced effector differentiation. Key questions remain about the tissue residency and memory capabilities of the Trim28-deficient TRM-like cluster.

Functional validation demonstrated Trim28-deficient TILs were more frequently TCF7-positive and, despite being underrepresented in tumors at baseline, increased their relative intratumoral recovery after anti-PD-1 treatment. This pattern suggests that Trim28 deficiency preserves a less differentiated pool with residual capacity to expand in response to checkpoint blockade.

However, because Trim28 loss was constitutive in these experiments, the same differentiation constraint that maintained these states likely continued to limit cytotoxic differentiation and intratumoral effector accumulation, preventing improved tumor growth control or survival. Thus, the Trim28 phenotype highlights that preserving stem-like potential is not sufficient for tumor clearance unless it can be coupled to appropriately timed effector differentiation.

Together, these observations highlight a therapeutic tension between effector differentiation and preservation of the progenitor-like pool. Rapidly growing transplantable mouse tumor models and short-term tumor-control endpoints are indispensable, but they can preferentially nominate perturbations that drive strong activation, clonal expansion, and cytotoxic differentiation as these features are often required for rapid tumor regression. However, persistent or repeated exposure to strong differentiating signals may erode the less differentiated progenitor cells needed for long-term persistence, proliferative renewal, and continued responsiveness to ICB.^2,3,5,8,34,39–41^

Our Trim28 experiments reveal the converse failure mode: constitutive inhibition of terminal cytotoxic differentiation may maintain less differentiated T cell states, but if differentiation remains blocked, these cells may fail to generate sufficient intratumoral cytotoxic effectors for tumor control. This distinction may be relevant to T cell-directed therapies more broadly, where strong T cell activation can produce initial anti-tumor responses but durability can be limited by exhaustion, reduced persistence, antigen escape, acquired resistance, or toxicity under persistent antigen exposure or continuous therapeutic stimulation.^42–45^ Thus, durable immunotherapy may benefit from temporal control of T cell fate rather than constitutive enforcement in either direction: perhaps alternating between waves of effector differentiation and periods of limited differentiation to preserve the renewable progenitor-like pool.^45,46^ Future experiments could further explore this idea with reversible methods of Trim28 deficiency or inhibition.

Several limitations of this manuscript should be considered. Clonal barcoding reduces false positives caused by expanded clones but also lowers sample size for clone-level tests. The differentiation-rate model is intentionally simplified and provides parameter ranges rather than direct biological rates. Finally, the TRM-like designation is based on transcriptional/accessibility signatures and technically challenging tissue distribution experiments; direct residency and memory-function assays remain needed.

## Methods

### Mice

All mouse experiments were performed in accordance with institutional guidelines and protocols approved by the Broad Institute IACUC under protocol #0054-05-15, “Characterization of genes regulating immunity.” Mice were maintained under specific pathogen-free conditions. Experimental mice were 6-14 weeks old. C57BL/6J mice were obtained from The Jackson Laboratory (JAX stock #000664). Donor and recipient strains included H11-Cas9 mice (B6J.129(Cg)-Igs2tm1.1(CAG-cas9*)Mmw/J, JAX stock #028239), OT-I mice (C57BL/6-Tg(TcraTcrb)1100Mjb/J, JAX stock #003831), congenic B6 CD45.1/Pep Boy mice (B6.SJL-Ptprca Pepcb/BoyJ, JAX stock #002014), and Tcf7-GFP reporter mice (B6(Cg)-Tcf7tm1Hhx/J, JAX stock #030909), crossed as needed to generate Cas9 x OT-I donors, reporter-bearing donors, and congenic Cas9 x CD45.1 or CD45.1/CD45.2 recipients, and P14 (Taconic B6.Cg-Tcratm1Mom Tg(TcrLCMV)327Sdz backcrossed 10 generations to Jackson C57BL/6J) TCR transgenic mice on the CD45.1 (B6.SJL-Ptprca Pepcb/BoyJ) congenic background.

### B16-OVA and Tumor Implantation

B16-OVA melanoma cells were B16.1 pLX311 Cas9 hygro-PGK-OVA cells obtained from the Manguso laboratory/Broad Institute and were confirmed mycoplasma negative before use. Cells were cultured under standard tissue-culture conditions in complete DMEM containing 10% FCS, 1% penicillin-streptomycin, and 1% GlutaMAX. For tumor implantation, cells were used while in log-phase growth, harvested, washed, resuspended in sterile PBS, and implanted subcutaneously into recipient mice. Tumors were measured with calipers, and tumor volume was calculated as length x width x width / 2. Mice were monitored and euthanized according to institutional protocol.

### Immune Checkpoint Blockade

For checkpoint-blockade experiments, mice received anti-PD-1 or isotype control antibodies by intraperitoneal injection at 200 ug per mouse per dose. Anti-PD-1 treatment used Bio X Cell anti-mouse PD-1 (CD279), clone RMP1-14 (catalog BE0146), and control treatment used Bio X Cell rat IgG2a isotype control, anti-trinitrophenol, clone 2A3 (catalog BE0089). In experiments where treatment was initiated based on tumor size, treatment began when tumors reached approximately 50 mm3.

### Naive CD8 T Cell Isolation

Naive CD8 T cells were isolated from spleens and lymph nodes of Cas9-expressing donor mice, typically H11-Cas9 x OT-I mice or related congenic combinations, using magnetic bead enrichment. Briefly, spleens and lymph nodes were processed into single-cell suspensions, red blood cells were lysed when necessary, and naive CD8 T cells were purified using a Miltenyi naive CD8 T cell isolation workflow. Purified cells were maintained in complete T cell medium supplemented with recombinant IL-7 and IL-15 at 10 ng/mL each. Cells were not activated through the TCR during the initial transduction period unless explicitly stated for in vitro activation or screening readouts.

### sgRNA Library Design and Cloning

sgRNAs for custom libraries were designed with Broad CRISPR design tools and selected to target candidate regulators nominated from earlier screens or from follow-up biological hypotheses. Pooled sgRNA oligonucleotides were cloned by Golden Gate assembly into pRDA_206_Thy1.1, transformed into Endura electrocompetent cells, and prepared using ZymoPURE maxiprep kits. Guide identities and 5’ to 3’ sequences for custom sgRNA libraries are listed in the guide-reference table (Table M1).

### Lentiviral Vectors, Production, and T Cell Transduction

Pooled sgRNA libraries were cloned into lentiviral vectors carrying an sgRNA cassette and a Thy1.1 surface marker to enable enrichment of transduced cells. For custom pooled libraries used in the in vivo single-cell and validation experiments, sgRNAs were cloned into pRDA_206_Thy1.1, a lentiviral sgRNA expression vector containing a U6-sgRNA cassette and a PGK-Thy1.1 transduction marker. Genome-wide screens used the BRIE whole-genome CRISPR knockout library cloned in pXPR_045 (Addgene plasmid #107144).

Lentivirus was produced in Lenti-X 293T or HEK293T cells by transient transfection of the sgRNA library plasmid with psPAX2 lentiviral packaging plasmid (Addgene #12260), pCMV-VSV-G envelope plasmid (Addgene #8454), and pCAG-Eco ecotropic MLV envelope plasmid (Addgene #35617). Viral production included both VSV-G and MMLV ecotropic envelope components to support infection of naive mouse T cells. The standard T175-scale production protocol used approximately 58 ug total DNA per flask: 28 ug transfer/library plasmid, 21 ug psPAX2, 4.5 ug pCMV-VSV-G, and 4.5 ug pCAG-Eco, transfected with Lipofectamine 3000/P3000 in Opti-MEM. Four hours after transfection, medium was replaced with viral harvest medium consisting of DMEM supplemented with 10% FBS, GlutaMAX, HEPES, MEM non-essential amino acids, penicillin-streptomycin, and BSA. Viral supernatant was collected approximately 48 hours after transfection, clarified by low-speed centrifugation, filtered through 0.45 um PES filters, aliquoted, and stored at −80 degrees C.

For T cell transduction, tissue-culture plates were coated with RetroNectin recombinant human fibronectin fragment (Takara, T100B), virus was bound to the plate surface by centrifugation, and viral harvest medium was removed before adding T cells. The standard protocol used RetroNectin at approximately 18 ug/cm2, followed by virus spin-loading at 2,000 x g for 2 hours at 32 degrees C. Naive CD8 T cells were added in complete T cell medium containing recombinant mouse IL-7, recombinant mouse IL-15, and LentiBOOST transduction enhancer. Cells were cultured on virus-coated plates for approximately 2 days and maintained without TCR activation through day 5 post-transduction. Viral input was titrated directly on primary T cells to select a dose that provided sufficient Thy1.1+ recovery while limiting excessive multi-integration. In pooled single-cell experiments, transduction was generally targeted to approximately 20-40% Thy1.1+ cells.

On day 5 after transduction, Thy1.1+ cells were enriched by staining with anti-Thy1.1-PE followed by anti-PE magnetic bead selection. Enriched cells were counted and used for in vitro screens, adoptive transfer, or input genomic DNA collection.

### Adoptive Transfer of T Cells into Mice

For in vivo Tcf7 sort screens, Perturb-seq screens, and validation experiments, Cas9-expressing OT-I CD8 T cells were transduced with pooled sgRNA libraries targeting candidate regulators and control guides, enriched for Thy1.1+ cells, and transferred into congenic recipient mice bearing or subsequently implanted with B16-OVA tumors. Donor and recipient populations were distinguished by congenic markers including CD45.1 and CD45.2, together with Thy1.1 and OT-I/TCR markers were used.

### Competitive Transfer Experiments

Naive CD8+ T cells were isolated from CD45.1 and CD45.1+/CD45.2+ mice. Donor and recipient populations were distinguished using congenic CD45.1 and CD45.2 markers. Recovered lymphocytes were gated on live Va2+ CD8+ T cells to identify OT-I CD8+ T cells, then gated on CD45.1/CD45.2 to distinguish control and experimental cells. Transduced experimental cells were identified by Thy1.1 expression.

### Tumor, Lymph Node, and Tissue Processing

Tumors were excised, minced, and enzymatically dissociated using Miltenyi tumor dissociation reagents or an equivalent tumor-infiltrating lymphocyte dissociation workflow. Dissociated tumor material was filtered to remove debris and enrich viable lymphocytes. Depending on experiment and sample quality, lymphocytes were enriched by Lympholyte-M density-gradient separation and/or magnetic enrichment before flow sorting. Draining lymph nodes were mechanically dissociated through a cell strainer to generate single-cell suspensions and processed separately from tumors.

Single-cell suspensions were stained with viability dye, Fc block, and surface antibodies to identify live CD8 T cells, donor-derived transferred cells, and transduced Thy1.1+ cells. For transcription-factor readouts, cells were fixed, permeabilized, and stained intracellularly for Tcf7, Prdm1, Eomes, Tox, and other markers using the eBioscience Foxp3/Transcription Factor Staining Buffer Set (Thermo Fisher/Invitrogen, catalog 00-5523-00) according to the manufacturer’s instructions. Flow-cytometry data were acquired on a Beckman Coulter CytoFLEX LX. Cell sorting into 96-well plates, including bulk RNA-seq sorting, was performed on a Sony cell sorter. For single-cell RNA-seq experiments, donor Thy1.1+ CD8 T cells were sorted before 10x loading using a Miltenyi cell sorter to maximize cell viability for single-cell capture.

For TotalSeq hashtag staining, purified or enriched cell suspensions from individual mice were stained with BioLegend TotalSeq-C mouse hashtag antibodies according to the manufacturer’s protocol, including repeated washes to reduce ambient hashtag oligonucleotide background. Hashtag-labeled samples were pooled for 10x library preparation.

### Guide-Count Library Preparation

Genomic DNA was isolated from sorted cell populations or tissue samples using QIAamp DNA Mini Kit/QIAamp Mini spin-column workflows, or by SPRI-based recovery for fixed or low-input sorted material where appropriate. For tumor-infiltrating T cell and tissue guide-count samples, tumor digests were enriched over Lympholyte-M and/or with CD8 TIL MicroBeads before pelleting or freezing for genomic DNA isolation where indicated by the experiment workflow.

Integrated sgRNA cassettes were amplified from genomic DNA using indexed Illumina-compatible primers following Broad/GPP pooled-screen PCR protocols. For standard pXPR guide-count PCR, up to 10 ug genomic DNA was amplified per 100 uL reaction using Titanium Taq DNA polymerase, DMSO, a staggered P5 primer mix, and a unique indexed P7 primer. In lower-input or nested tissue workflows, PCR cycle number was selected by qPCR using Q5 Hot Start high-fidelity polymerase, and samples were split across multiple reactions to use available genomic DNA while limiting PCR jackpotting. No-template controls and test gels were included to check for contamination and expected amplicon size.

PCR products were pooled by sample and purified by AMPure XP cleanup, double-sided AMPure XP size selection (0.6x/1.2x), or gel extraction depending on amplicon workflow and input complexity. Purified amplicons were quantified by Qubit and Bioanalyzer/TapeStation, pooled, and sequenced on Illumina instruments. Guide counts were generated with PoolQ^47^ using sample-barcode or condition tables and the corresponding guide sequence sets listed in Table M1 for custom libraries or the BRIE whole-genome reference for the genome-wide screen.

### Bulk RNA-seq Library Preparation

Bulk RNA-seq FASTQ files were quantified with Salmon v1.4.0 against the GENCODE vM25 mouse transcriptome using automatic library-type detection, selective alignment (--validateMappings), and GC-bias correction (--gcBias).^48,49^ Transcript-level Salmon NumReads values were summed to gene-level counts, rounded to integers, and filtered to retain genes with nonzero total counts. Gene-level counts were normalized with DESeq2 median-ratio size factors.^50^ Variance-stabilized expression was computed with vst(blind=TRUE) for PCA, and DESeq2-normalized counts were used for heatmaps and correlation analyses.

### Validation of Naive T Cell Transduction in the LCMV Model

To compare naive-preserving lentiviral transduction with activation-dependent transduction in vivo, P14 LCMV-specific CD8 T cells were prepared under two pre-transfer conditions. For the lenti naive condition, naive P14 CD8 T cells were transduced using the naive T cell transduction workflow described above and maintained without TCR stimulation before transfer. For the lenti-activated condition, P14 CD8 T cells were transduced and activated with anti-CD3/CD28 stimulation before transfer. Each experimental population was mixed with freshly isolated congenically marked naive P14 CD45.1+CD45.2+ CD8 T cells as cotransferred controls. Recipient mice were infected with LCMV Armstrong after transfer, and splenic P14 cells were analyzed 7 and 30 days after infection.

Pre-transfer RNA-seq comparisons included freshly isolated naive P14 cells, lenti naive P14 cells, and anti-CD3/CD28-activated non-transduced comparator cells. Day-7 RNA-seq comparisons included recovered P14 cells derived from lenti naive or lenti-activated donor cells and from matched cotransferred naive controls. At day 30, central memory and effector memory P14 CD8 T cells were flow sorted from control and experimental populations before bulk Smart-seq2 library preparation. Flow-cytometry analyses quantified recovery of experimental and control P14 populations, short-lived effector cells at day 7, and central memory cells at day 30.

### Genome-Wide and Focused In Vitro CRISPR Screens

Genome-wide and focused in vitro CRISPR screens were performed in Cas9-expressing naive CD8 T cells using the lentiviral transduction workflow described above. After transduction, cells were cultured under acute- or chronic-stimulation conditions, with a separate hypoxic condition for the Tox screen. For acute stimulation, purified transduced T cells were stimulated overnight on anti-CD3/CD28-coated plates, then removed from stimulation plates and cultured in cytokine-containing T cell medium. For chronic stimulation, cells remained on anti-CD3/CD28-coated plates throughout the experiment. For hypoxia screens, cells were cultured under 1% oxygen. Anti-CD3 and anti-CD28 coating used clones 145-2C11 and 37.51 at 5 ug/mL each for Tcf7 in vitro screen conditions.

Cells were sorted into transcription-factor high and low populations for Tcf7, Prdm1, Eomes, or Tox, and guide abundance was quantified in each sorted population. Tcf7, Prdm1, Eomes, and Tox screens used antibody staining to identify high- and low-expressing cells, except where Tcf7-GFP reporter sorting was used.

For the BRIE genome-wide screen, naive CD8 T cells were isolated from 28 Cas9 x OT-I mice, yielding approximately 200 million T cells, and transduced with the BRIE whole-genome CRISPR knockout library. Five days after transduction, Thy1.1+ cells were magnetically purified. Fifty million purified cells were resuspended in T cell medium supplemented with 10 ng/mL IL-7, 10 ng/mL IL-15, and 2 ng/mL IL-2, then stimulated overnight on plates coated with anti-CD3 and anti-CD28 antibodies. Cells were then lifted and resuspended in T cell medium supplemented with IL-7, IL-15, and IL-2 with folate concentration adjusted to high or low conditions. Fifty million cells per condition were collected on day 3 for genomic DNA isolation. An additional 50 million cells per condition were restimulated overnight on anti-CD3/CD28-coated plates, lifted, and resuspended in the same folate condition. Cells were maintained in culture until day 7 from initial activation, at which point genomic DNA was isolated from recovered cells, sgRNA cassettes were amplified by indexed PCR, and libraries were sequenced on an Illumina NextSeq using custom sequencing primers.

### Processing and Statistical Analysis of in vitro Screens

Guide sequencing reads were processed with PoolQ^47^. For each screen comparison, PoolQ reported guide-level log2 fold changes from log-normalized counts and gene-level ranked-list enrichment scores that quantify whether guides targeting a gene were concentrated at either tail of the guide-level log2 fold-change distribution. In Figure 1e-f and Figure S2a, we report the PoolQ average guide-level log2 fold change for each gene and the average -log10(P) from the more significant ranked-list direction.

### In Vivo Tcf7 Sort CRISPR Screens

For in vivo Tcf7 sort screens, Cas9-expressing OT-I CD8 T cells were transduced with pooled sgRNA libraries targeting candidate Tcf7 regulators and control guides, enriched for Thy1.1+ cells on day 5 after transduction, and transferred into congenic recipient mice. One day after transfer, recipients were implanted bilaterally with 1 × 10^6 B16-OVA cells per flank. Transferred cells were recovered from tumors and draining lymph nodes after in vivo differentiation, typically 14 days after tumor implantation, and sorted into Tcf7-high and Tcf7-low populations by Tcf7-GFP reporter expression or Tcf7 antibody staining, as dictated by the experiment. Genomic DNA from sorted Tcf7-high and Tcf7-low populations was isolated, sgRNA cassettes were amplified with indexed primers, and amplicons were sequenced.

The in vivo Tcf7 sort-screen libraries targeted follow-up genes selected from the whole-genome Tcf7 screen, with two sgRNAs per gene and non-targeting and intergenic controls. The first library contained 198 targeting sgRNAs across 99 genes, 33 NO_SITE controls, and 27 intergenic controls; the second contained 200 targeting sgRNAs across 100 genes, 32 NO_SITE controls, and 25 intergenic controls (Table M1). These libraries did not include essential-gene controls.

Across the two B16-OVA in vivo Tcf7 sort screens, input cells, draining-lymph-node samples, tumor-infiltrating Tcf7-high and Tcf7-low fractions, and plasmid library controls were processed for guide-count sequencing. Where relevant, male and female samples were processed as separate input or lymph-node samples before guide-level analysis.

### 10-Gene Perturb-seq Screen

The 10-gene Perturb-seq experiment was designed to profile how candidate regulators nominated from the in vitro and in vivo Tcf7 screens altered CD8 T cell state, clonal expansion, and response to checkpoint blockade in B16-OVA tumors. The sequenced single-cell dataset contains 7 isotype-treated and 9 anti-PD-1-treated recipient mice.

The 10-gene Perturb-seq library targeted Setdb1, Trim28, Ifngr1, Ifngr2, Jak2, Jak3, Tnfrsf1b, Il18r1, Prdm1, and Bach2, with two sgRNAs per gene. The library also contained non-targeting and intergenic control sgRNAs, including NO_SITE and ONE_INTERGENIC_SITE controls (Table M1).

Naive CD8 T cells were isolated from 12 Cas9 x OT-I donor mice and transduced with the 10-gene Perturb-seq sgRNA library on RetroNectin-coated plates. On day 5 after transduction, approximately 20% of cells were Thy1.1+. Thy1.1+ cells were enriched by Thy1.1-PE staining followed by anti-PE magnetic bead selection, yielding approximately 6 × 10^6 cells. A post-enrichment flow check showed 83% Thy1.1+ cells. Approximately 230,000 enriched cells were transferred per Cas9 x CD45.1 recipient mouse. One day after transfer, recipients were implanted bilaterally with subcutaneous B16-OVA tumors at 1 × 10^6 cells per flank.

Tumors were measured before checkpoint-blockade assignment, and mice were assigned to anti-PD-1 or isotype treatment to balance tumor volume across treatment groups. Treatment began 10 days after tumor implantation. Two mice were euthanized at treatment initiation because of ulcerated tumors. The remaining 16 mice were harvested 5 days after treatment initiation. For each mouse, both tumors were pooled into one Miltenyi C tube containing RPMI plus tumor-dissociation enzymes, minced, processed with the gentleMACS 37C m_TDK1 program, filtered through 70 um strainers, enriched over Lympholyte-M, and then enriched with the Miltenyi CD8 TIL MicroBeads workflow. Cells were blocked and stained with viability dye, Fc block, TotalSeq hashtag antibodies, CD45.1, CD45.2, and Thy1.1. Isotype-treated and anti-PD-1-treated cells were pooled separately before sorting. The final sorted pools contained approximately 132,000 isotype cells and 200,000 anti-PD-1 cells and were loaded across eight 10x 5’ v2 channels.

Single-cell libraries were prepared using the 10x Genomics Chromium 5’ workflow with gene-expression, feature-barcode/hashtag, and direct CRISPR guide-capture libraries, using an RT direct sgRNA-capture oligo spike-in. Approximately 40,000-50,000 cells were targeted per 10x channel.

### Trim28/Setdb1 Perturb-seq Screen

Trim28/Setdb1 Perturb-seq was performed as a focused clonally barcoded single-cell CRISPR screen in B16-OVA tumors.

The Trim28/Setdb1 Perturb-seq library contained ten sgRNAs targeting Trim28, ten sgRNAs targeting Setdb1, and six NO_SITE control sgRNAs. Trim28-1, Trim28-2, Setdb1-1, and Setdb1-2 were the same core guide identities used in the 10-gene Perturb-seq screen; library-specific sequences are listed in Table M1.

Naive CD8 T cells were isolated from 18 Cas9 x OT-I donor mice and transduced with the Trim28/Setdb1 Perturb-seq library. Thy1.1+ cells were enriched on day 5 after transduction. Recipient mice included congenic mice from Cas9 x CD45.1 and Cas9 x CD45.1/CD45.2 backgrounds. Recipients were implanted bilaterally with 4 × 10^5 B16-OVA cells per flank. Transduced donor T cells were transferred 1 day after tumor implantation, with a target dose of 2 × 10^5 Thy1.1+ cells per mouse; approximately 1.7 × 10^5 Thy1.1+ cells were transferred per mouse for most mice.

Mice were treated with anti-PD-1 or isotype control after tumors reached approximately 50 mm3, 11 days after tumor implantation. Mice were harvested in two batches 6 and 11 days after treatment initiation. At harvest, tumors and draining lymph nodes were collected. Tumors were dissociated to single-cell suspensions and enriched for donor T cells before sorting and 10x loading. Draining lymph nodes were processed separately. Cells from each mouse were hashtagged, pooled by tissue and harvest batch, and processed using the 10x Genomics Chromium 5’ workflow with gene-expression, CRISPR guide-capture, and feature-barcode libraries.

### Trim28 Multiome Screen

Naive Cas9-expressing OT-I CD8 T cells were transduced and Thy1.1-enriched as described above. NT control and Trim28-1 gRNA-transduced cells were generated in congenically distinguishable donor populations, mixed at approximately equal ratios, and transferred by tail-vein injection into recipient mice. One day after transfer, mice were implanted bilaterally with 2.5 × 10^5^ B16-OVA cells per flank. Tumors were harvested approximately two weeks after implantation, processed to single-cell suspensions, enriched for CD8 TILs, and stained for CD8, Thy1.1, CD45.1, and CD45.2 as described above. Donor Thy1.1-positive NT control and Trim28-1 cells were sorted as separate populations based on congenic markers.

Sorted cells were processed immediately using the 10x Genomics Chromium Next GEM Single Cell Multiome ATAC + Gene Expression workflow. Nuclei were isolated, counted, and loaded onto the Chromium controller, with approximately 5,000 nuclei loaded based on the pre-loading count. Transposition, GEM generation and barcoding, cDNA amplification, and construction of paired gene-expression and ATAC libraries were performed according to the manufacturer’s protocol. Condition-specific GEX libraries were pooled separately from condition-specific ATAC libraries and sequenced by Broad Walk-Up Sequencing on an Illumina NextSeq 2000 P2. The ATAC pool was sequenced with paired-end 50/49 bp reads and 8 bp/24 bp index reads, and the GEX pool was sequenced with paired-end 28/90 bp reads and 10 bp dual-index reads.

### Trim28 Validation and Competitive-Transfer Experiments

For Trim28 validation experiments, naive Cas9-expressing OT-I CD8 T cells were transduced with NT, Trim28-1, Trim28-2, or mixed NT/Trim28 sgRNA constructs and enriched for Thy1.1+ cells before transfer into B16-OVA-bearing recipient mice. Competitive-transfer experiments mixed NT and Trim28-perturbed T cells at defined input ratios before transfer. Input ratios were measured by flow cytometry or guide counts, and recovered ratios were measured in draining lymph nodes, tumors/TIL, skin, or other tissues at harvest.

For flow-based Tcf7 validation, recovered donor cells were identified by congenic and Thy1.1 markers, and Tcf7 expression was quantified in NT versus Trim28-perturbed cells. For guide-count validation, genomic DNA was isolated from sorted T cells or tissue samples, and NT and Trim28 sgRNA representation was quantified by amplicon sequencing and PoolQ. Trim28 recovery in tumors was normalized to input or to matched draining lymph node samples as indicated in the figure legends. For anti-PD-1 competitive-transfer experiments, mice received isotype control or anti-PD-1 treatment after tumors reached the experiment-specific treatment threshold, typically approximately 50 mm3. For therapeutic tumor-growth experiments, mice received OT-I T cells transduced with NT or Trim28-1 sgRNAs, followed by B16-OVA challenge and tumor monitoring.

### Perturb-seq Processing

#### scRNA-seq data pre-processing

For the 10-gene Perturb-seq experiment, gene expression unique molecular identifier (UMI) counts and HTO counts were generated with Cell Ranger v7.0.1 using the mm10 reference.^51^ Cells were filtered out if more than 20% of reads mapped to mitochondrial genes. We removed TCR genes. We assigned mouse hashtags using HTODemux in Seurat.^52,53^ We filtered for doublets by keeping only cells assigned to a single hashtag. Treatment in the 10-gene Perturb-seq experiment was defined by 10x lane, and mouse identity was defined by the assigned hashtag within that treatment arm. Lanes 1-4 contained cells from isotype-treated mice labeled with hashtags 1-7; lanes 5-8 contained cells from anti-PD-1-treated mice labeled with hashtags 1-8, with hashtag 8 representing two anti-PD-1-treated mice. For the Trim28/Setdb1 Perturb-seq experiment, each hashtag corresponded to one mouse. Treatment condition was assigned from the mouse hashtag key. Hashtags 1-12 were used in the first harvest, and hashtags 13-24 were used in the second harvest. Lanes 1 and 9 were lymph-node libraries, and the remaining lanes were tumor libraries. Next, we assigned clonal barcode and gRNA constructs to cells (Clonal Barcode Processing Section). During clonal barcode processing, we removed cells that were doublets based on construct composition. We kept cells without guides.

The single-cell RNA sequencing data were processed using the Scanpy toolkit.^54^ Normalization was performed using total counts normalization (sc.pp.normalize_total) with the exclusion of highly expressed genes to mitigate bias. Subsequently, a logarithmic transformation (sc.pp.log1p) was applied. Highly variable genes were identified using the Seurat method^53^, retaining the top 8,000 genes. The data was scaled to unit variance and clipped at a maximum value of 10. Using the top 50 principal components, we constructed a smoothed k-nearest neighbor graph (k=20), generated a UMAP embedding^55^, and performed Leiden clustering at multiple resolutions^56^. In both the 10-gene Perturb-seq and Trim28/Setdb1 Perturb-seq experiments, a pre-filter high-mito population was excluded during quality-control filtering before downstream analyses. We also calculated a z-score normalized layer using only control (NT) cells as reference, where the mean and standard deviation were computed from non-targeting control cells only and then applied to all cells in the dataset. This control-normalized z-score layer was used for the dotplots and gene set scoring.

### Construct Processing

#### Clonal Barcode and Guide RNA Extraction

Guide/construct libraries were sequenced as paired-end FASTQs: the forward read carried the cell barcode and molecular UMI, while the reverse read carried the guide sequence and clonal barcode flanked by known adapters. We used Pycashier^57^ to leverage the adapter context and extract clonal barcodes and guide sequences. To assign guide identities, guide read sequences were aligned with STAR^58^ to the corresponding 10-gene Perturb-seq or Trim28/Setdb1 Perturb-seq sequence set (Table M1). Reads were deduplicated by molecular UMI, cell barcode, clonal barcode, and guide. We then constructed a cell-barcode–by-construct count table in which each construct is a unique guide–clonal barcode pair and each entry is the number of deduplicated molecules observed for that construct in that cell.

#### Assigning Constructs to Cells

From this count matrix, we retained only cell-barcode–construct entries above a minimum threshold (10-gene Perturb-seq: >10; Trim28/Setdb1 Perturb-seq: >3), yielding a binary cell-barcode-by-construct matrix. We chose these thresholds empirically from single-construct clones with many cells, which we treated as bona fide correctly assigned cells because many cells shared the same construct. In the 10-gene Perturb-seq dataset, the large single-construct clones had construct counts extending close to the retained cutoff, down to 11 molecules (Figure S4a). In the Trim28/Setdb1 Perturb-seq dataset, analogous large single-construct clones extended down to 4 molecules. Within these clones, construct-count distributions were often approximately unimodal. This pattern suggests that the effective detection threshold is frequently clone-specific, possibly reflecting clone-specific lentiviral integration context, rather than a consistent shift by mouse, treatment, or guide identity. We therefore used a permissive dataset-level cutoff to assign constructs to clones, then relied on the linear-program filter to remove cells with incompatible construct assignments.

This differs from the default Cell Ranger CRISPR guide-calling procedure (10x Genomics Cell Ranger CRISPR Guide Capture algorithm^51^), which fits a two-component Gaussian mixture separately for each gRNA on log-transformed guide UMI counts and derives a gRNA-specific threshold.

For each cell we encoded the set of retained constructs and defined a putative clone as a unique construct combination.

We observed construct sharing patterns consistent with doublets: many cells carried construct A, many carried construct B, and a small number carried both A and B (Figure S4b). To resolve construct sharing between putative clones, we formulated a linear program that selects putative clones to maximize the number of cells retained, under the constraint that each construct is assigned to at most one clone. Let i index putative clones, let n_i be the number of cells with clone i, and let x_i indicate whether clone i is retained. Define a_ij = 1 if construct j appears in clone i. We solved:

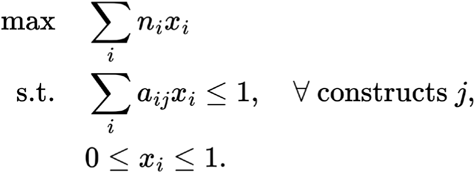

We solved the LP with HiGHS^59^ and retained clones with x_i = 1. We assume that clones that were not chosen by the LP are doublets.

To verify that clonal barcode could be used for doublet calling, we compared clonal doublet assignments with mouse hashtag oligo doublet calling in the 10-gene Perturb-seq dataset (Figure S4c). The two assays capture different subsets of doublets: mouse hashtag oligo calling misses doublets formed from two cells from the same mouse, whereas clonal doublet calling requires retained construct evidence from both contributing cells. Despite this discrepancy, 52.1% of mouse hashtag oligo doublets were called clonal doublets, compared with 7.6% of mouse hashtag oligo singlets, indicating strong association of the two doublet calling approaches (Figure S4c; P < 1e-300).

### Perturb-seq Analysis

#### Determining Perturbation for Cells

In the 10-gene Perturb-seq experiment, cells labeled as NT or as a given perturbation were restricted to cells with exactly one retained guide assignment. For the Trim28/Setdb1 Perturb-seq dataset, we instead collapsed multi-guide cells to curated gene-level labels. Cells containing Setdb1 and no Trim28 guide were labeled Setdb1, including cells that also carried control (NO_SITE) guides or multiple Setdb1 constructs; cells containing Trim28 and no Setdb1 guide were labeled Trim28; cells containing both genes were labeled Setdb1+Trim28; and cells carrying only control guides were labeled NT.

#### Differential Gene Expression Analysis

DEG analysis was performed between a query set of cells and a background set of cells. The query cells were either a cell type or perturbation and the background was either all other cell types or NT cells respectively. We used the two-sided Wilcoxon rank-sum test on normalized gene expression values. The P values obtained from the tests were then adjusted using the Benjamini-Hochberg adjustment. Genes were tested only if at least 5% of query cells had expression of the gene. The log2 fold change (logFC) is calculated as log2((mean expression in group1 + 0.01) / (mean expression in group2 + 0.01)).

#### Gene Set Scoring

A gene set signature score for a cell is derived by computing the average log normalized z-score expression of a predefined list of genes and subtracting the global average log normalized z-score across all genes.

Differential expression–based progenitor and dysfunction signatures were obtained from Pritykin et al.^60^ Briefly, the authors assembled a compendium of 136 bulk RNA-seq CD8⁺ T cell datasets from mouse models of infection and cancer. Datasets were clustered by gene expression and annotated by state. We used the progenitor vs. dysfunctional differential expression analysis, selecting significant genes and retaining the top 20 with the highest and lowest logFC.

Progenitor vs. terminal exhaustion signatures were derived from Miller et al.^2^, who performed bulk RNA-seq of progenitor and terminal exhausted tumor-infiltrating CD8⁺ T cells. We selected significant differentially expressed genes and retained the top 20 upregulated and downregulated.

Core TRM signatures were obtained from Mackay et al.^19^, who performed whole-transcriptome microarray profiling of circulating memory and TRM cells from skin, lung, and gut after local viral infection. Significant genes upregulated in TRM across all three tissues were retained.

TRM transcriptional profiles were obtained from Mackay et al.^20^ The authors performed RNA-seq of antigen-specific CD8⁺ T cells from skin (HSV) and small intestine (LCMV), comparing TRM to central and effector memory subsets. From Mackay et al. Supplementary Figure 13a, we selected genes consistently upregulated in TRM across both tissues.

A pan-tissue TRM transcriptional program was derived from Milner et al.^17^ They integrated microarray datasets of antigen-specific TRM from small intestine, kidney, skin, lung, and brain across multiple viral infections. Differential expression was performed against splenic central and effector memory subsets, and genes consistently upregulated across all five TRM populations defined the core signature.

TRM residency signatures were obtained from Crowl et al.^18^ The authors profiled IV⁻ CD8⁺ P14 TRM cells from small intestine, kidney, liver, salivary gland, and adipose tissue after acute LCMV infection, comparing them to splenic memory cells by bulk RNA-seq. Differential expression and clustering across tissues identified genes consistently enriched in TRM.

Human pan-cancer TRM and TEX signatures were derived by Burn et al.^21^ The authors profiled CD8⁺ T cells by scRNA-seq from breast cancer tumours with matched non-cancerous breast tissue and from colorectal liver metastases with paired non-cancerous and donor liver tissue.

Within each dataset, CD103⁺ resident clusters were identified: clusters enriched in non-tumour tissues were defined as TRM, while those enriched in tumours were defined as TEX. Differential expression yielded TRM and TEX signatures, which were refined by focusing on leading-edge genes concordantly enriched across breast and liver to generate a pan-cancer TRM and TEX.

TRM and TR-TEX signatures were obtained from Park et al.^22^ Using single-cell TEA-seq, the authors profiled P14 CD8⁺ T cells in small intestine epithelium, salivary gland, and liver after acute (Armstrong, Arm) versus chronic (Clone 13, Cl13) LCMV 30 days post infection. They defined Arm TRM and Cl13 TR-TEX states, then derived cell-state–specific signatures by keeping genes upregulated in that state across ≥2 tissues relative to both matched cells of the other state and splenic circulating memory, excluding genes also increased in the other state vs circulating memory.

#### Testing Deviations from Expected Guide Abundance Cell/Clone Count Relationships

We built a null model using control guides to link one guide-abundance measurement to another recovered count. For each control guide, counts were modeled as clone/cell ∼ NB(μ = α·concentration, dispersion = β), with Var = μ + β·μ² (β = 0 reduces to Poisson). The parameter α is the linear scaling factor (doubling the x-axis abundance doubles the expected recovered count), and β captures overdispersion. We estimated α and β by maximum likelihood on control guides. For each guide, we computed the expected mean μ̂ = α̂·concentration, then used the fitted NB(μ̂, β̂) to assess deviation of the observed count. Two-sided p-values were computed by doubling the smaller of the lower-tail and upper-tail NB probabilities. Multiple-testing correction was performed via Benjamini–Hochberg FDR. The plotted band shows the 2.5% and 97.5% predictive quantiles for the fitted NB at each x-axis abundance (Figures 1h; 4a,b; 5b).

For the two in vivo Tcf7 sort screens shown in Figure 1h, we used a modified version of this model to stabilize the fit at zero recovered counts. Specifically, we added a pseudocount of 1 to both Tcf7- and Tcf7+ guide counts before fitting the control-guide relationship, such that for control guide i with Tcf7- count x_i and Tcf7+ count y_i, we fit y_i + 1 ∼ NB(mu_i, β) with mu_i = α(x_i + 1) and Var(y_i + 1) = mu_i + β mu_i^2. Guide-level deviations and two-sided p-values were then computed under this pseudocount-adjusted model. The two guide-level P values for each gene were converted to signed normal-equivalent Z-scores and combined using a signed Stouffer method to generate the gene-level summaries (Table S1.6; Table S1.7).

#### Clonal Modeling

We used a discrete-time branching process to estimate plausible ranges of progenitor division parameters from clone-level data. Each simulated clone began with one progenitor. The number of division cycles was drawn from a Poisson distribution with mean 5, chosen to match the observed clone size distribution. At each cycle, differentiated cells divided into two differentiated cells. Each progenitor underwent symmetric self-renewal with probability p_sym, asymmetric division with probability p_asym, symmetric differentiation with probability p_dif, or no division with the remaining probability 1 - (p_sym + p_asym + p_dif). After all divisions, each cell was retained independently to model incomplete capture. We used a retention probability of 0.15 for the 10-gene Perturb-seq experiment and 0.10 for the Trim28/Setdb1 Perturb-seq dataset.

These values were chosen from rough recovery estimates comparing the number of donor-derived cells expected in the mouse with the number of cells profiled after single-cell processing. For the 10-gene Perturb-seq experiment, we estimated that approximately 400,000 cells were donor-derived and profiled 68,000 cells, corresponding to an effective recovery fraction of approximately 0.17, and therefore used a nearby fixed value of 0.15. For the Trim28/Setdb1 Perturb-seq experiment, donor and endogenous cells were more difficult to distinguish in this experiment, so we assumed a lower effective recovery and used 0.10.

For each perturbation, we summarized observed and simulated clones as counts of progenitor and differentiated cells. We classified Progenitor states as those annotated Progenitor and Progenitor Cycling; all other states were treated as differentiated.

We compared observed and simulated clones using two summary statistics. First, we computed the empirical CDF of log2(clone size). Second, we fit a monotone-decreasing isotonic regression of progenitor fraction versus log2(clone size). We evaluated both curves on 100 equally spaced points from 0 to 0.9 times the maximum observed log2 clone size and defined loss as the sum of the mean-squared errors for the ECDF and isotonic curves (Figure 4g).

We evaluated loss on a grid of p_sym in [0, 0.5], p_asym in [0, 0.4], and p_dif in [0, 0.4], excluding points with p_sym + p_asym + p_dif > 1. We used 30 grid points per axis and simulated 1,000,000 surviving clones for each grid point. We summarized parameter ranges using the 100 lowest-loss points. To compare NT and Bach2, we computed the centroid of the 100 lowest-loss points for each perturbation, centered the two point clouds separately, and fit the first principal component to the pooled centered coordinates to obtain a shared direction. Each perturbation-specific line was then defined by its centroid and this shared direction. We obtained parameter ranges by intersecting each line with the constraints 0 ≤ p_sym, p_asym, p_dif ≤ 1 and p_sym + p_asym + p_dif ≤ 1.

### Multiome Processing

#### Multiome data pre-processing

We processed single-cell multiome (combined scRNA-seq + scATAC-seq) data with Cell Ranger ARC v2.0.2^61^ against the mm10 reference. Briefly, for each sample we obtained raw fragment files containing Tn5-transposase insertion sites. We then excluded low-quality cells based on total fragment counts (e.g., retaining cells with ≥100 unique insertions). We then applied a Tn5 offset (shifting the 5′ and 3′ ends by ±4–5 bp) to more accurately account for Tn5 insertion.

We first called peaks using MACS2^62^ on our single-cell ATAC data. We then merged these MACS2-called peaks with a curated set of CD8+ T cell peaks from A unified atlas of CD8 T cell dysfunctional states in cancer and infection.^60^ Because single-cell ATAC-seq can be sparse, incorporating this curated atlas helps ensure better coverage of biologically relevant CD8+ T cell specific regions.

Using Scanpy^54^, we constructed cell-by-peak count matrices by counting the number of Tn5 insertions falling within the intervals for each unified peak atlas. We performed library-size normalization by scaling each cell’s total insertion counts to a fixed value followed by a logarithmic transformation (sc.pp.log1p). We scaled the data to unit variance while clipping outliers at a maximum value of 10. We next computed 30 principal components, applied Harmony^28^ for batch correction between Control and Trim28 cells, and built a k-nearest neighbor graph (k=20). We then generated a UMAP embedding^55^, performed Leiden clustering^56^ at multiple resolutions, and annotated cells based on the expression of genes from the literature and our Perturb-seq screens.

#### Repetitive Element Analysis

We downloaded repeat regions annotated by RepeatMasker^63^ from the UCSC Genome Browser^64^ and retained only regions larger than 50 bp. For RNA, we used BAM files generated by Cell Ranger to count uniquely mapped reads fully contained within these repeat regions. For ATAC, we used the fragments file from Cell Ranger ARC to count insertion events within repeat regions. Counts were then aggregated across all regions sharing the same repeat name, resulting in a repeat name count matrix for each modality.

To validate the multiome RNA and ATAC counting procedures (Figure S15), we compared reconstructed counts from the raw BAM files with counts derived from Cell Ranger in the final processed multiome AnnData objects. We randomly sampled 100 retained genes from the multiome RNA dataset and 100 retained peaks from the multiome ATAC dataset. For RNA, we used the Cell Ranger gene-expression BAM files and counted unique cell-barcode/UMI pairs for uniquely mapped reads that were fully contained within GENCODE vM10^49^ gene intervals. For ATAC, we used the Cell Ranger ARC ATAC BAM files, applied the standard Tn5 offset to each read end, and counted insertion events falling within each sampled peak. For each sampled gene or peak, we then computed the Pearson correlation across cells between the reconstructed counts and the corresponding counts in the processed multiome AnnData object.

For library size normalization, we used a custom procedure that adjusts repeat counts for each cell based on sequencing depth. For RNA, the library size was calculated as the total counts in gene regions; for ATAC, it was the total counts in peak regions. We normalized repeat counts by dividing each cell’s repeat counts by its library size, then applied a global scaling factor to ensure the median library size of the normalized repeat count matrix matched the median library size of the original (unnormalized) repeat count matrix.

## Data and code availability

Raw and processed sequencing data generated in this study have been deposited in the Gene Expression Omnibus under accession numbers GSE334845 for bulk RNA-seq, GSE334846 for pooled bulk CRISPR screens, GSE334877 for the 10-gene Perturb-seq experiment, GSE334890 for the Trim28/Setdb1 Perturb-seq experiment, and GSE334878 for the Trim28 single-cell multiome experiment.

All code used to produce the analysis presented in the paper is available at https://github.com/pritykinlab/tcf7_clonal_perturbseq.

## Supplementary figure legends

**Figure S1.**
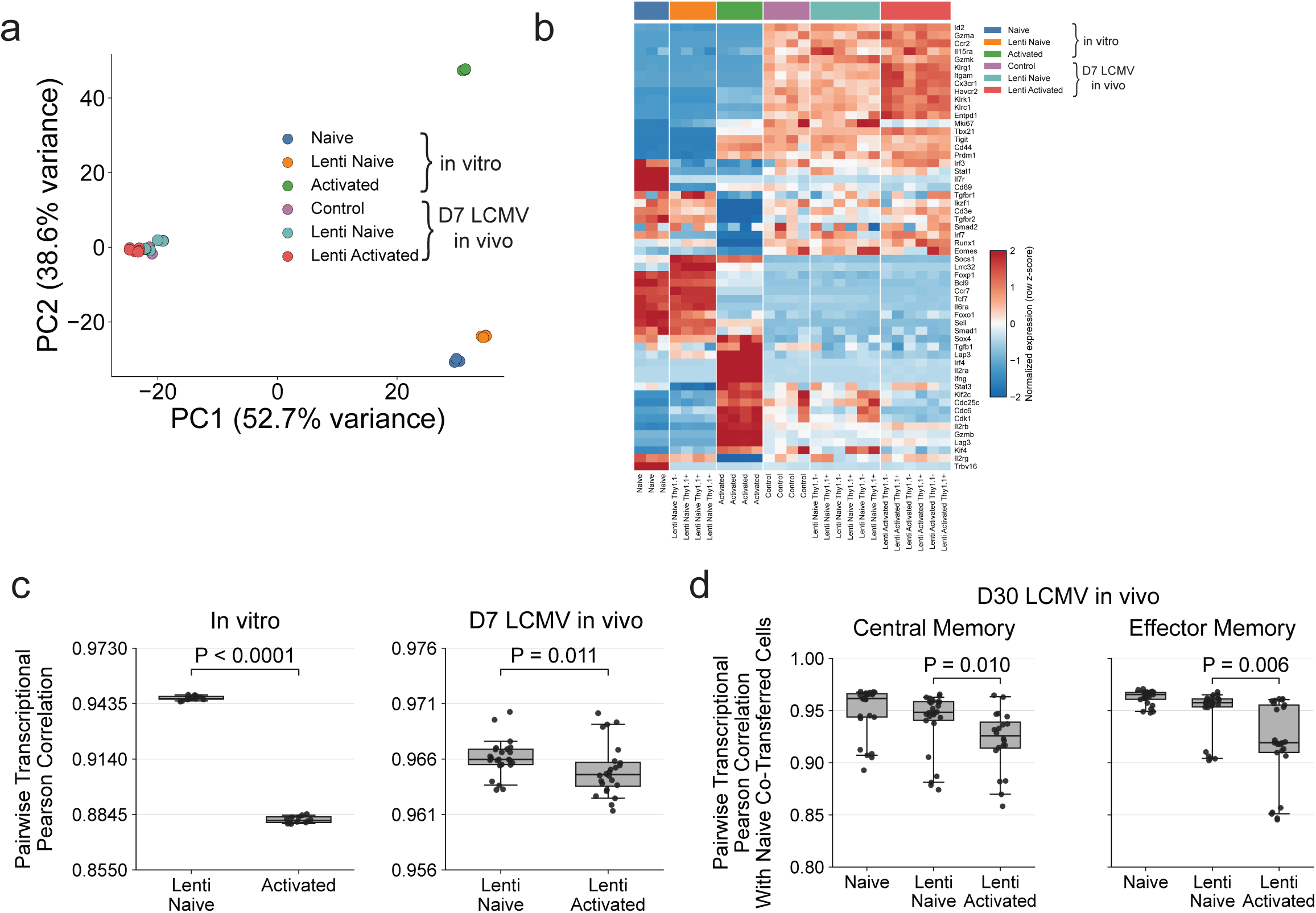
Additional analysis of *in vitro* and *in vivo* Tcf7 regulator screens. (a) Principal component analysis of DESeq2 variance-stabilized bulk RNA-seq profiles for the 27 RNA-seq samples underlying Figure 1c. (b) Heatmap showing DESeq2-normalized expression of the 27 individual RNA-seq samples from Figure 1c. Values are row z-scores computed across samples. Thy1.1 is the lentiviral transduction marker. (c) Pairwise Pearson correlations computed from log2-normalized gene expression for the samples shown in Figure 1c. Left, pre-transfer lenti naive and anti-CD3/CD28-activated samples were correlated with freshly isolated naive samples. Right, day-7 cells derived from lenti naive or lenti-activated P14 cells after LCMV infection were correlated with cells derived from cotransferred naive controls. Each point is one treatment-control sample-pair correlation. P values were calculated by treating pairwise sample correlations as observations and comparing the two displayed distributions within each panel using two-sided Mann-Whitney U tests. (d) Pairwise Pearson correlations between day-30 memory RNA-seq samples derived from control and cotransferred naive or transduced populations, split by flow-sorted central memory (CD44+CD62L+) and effector memory (CD44+CD62L-) P14 CD8 T cells. Each point is one control sample-pair correlation within the indicated input context. P values were calculated by treating pairwise sample correlations as observations and comparing lenti naive and lenti-activated distributions using two-sided Mann-Whitney U tests.

**Figure S2.**
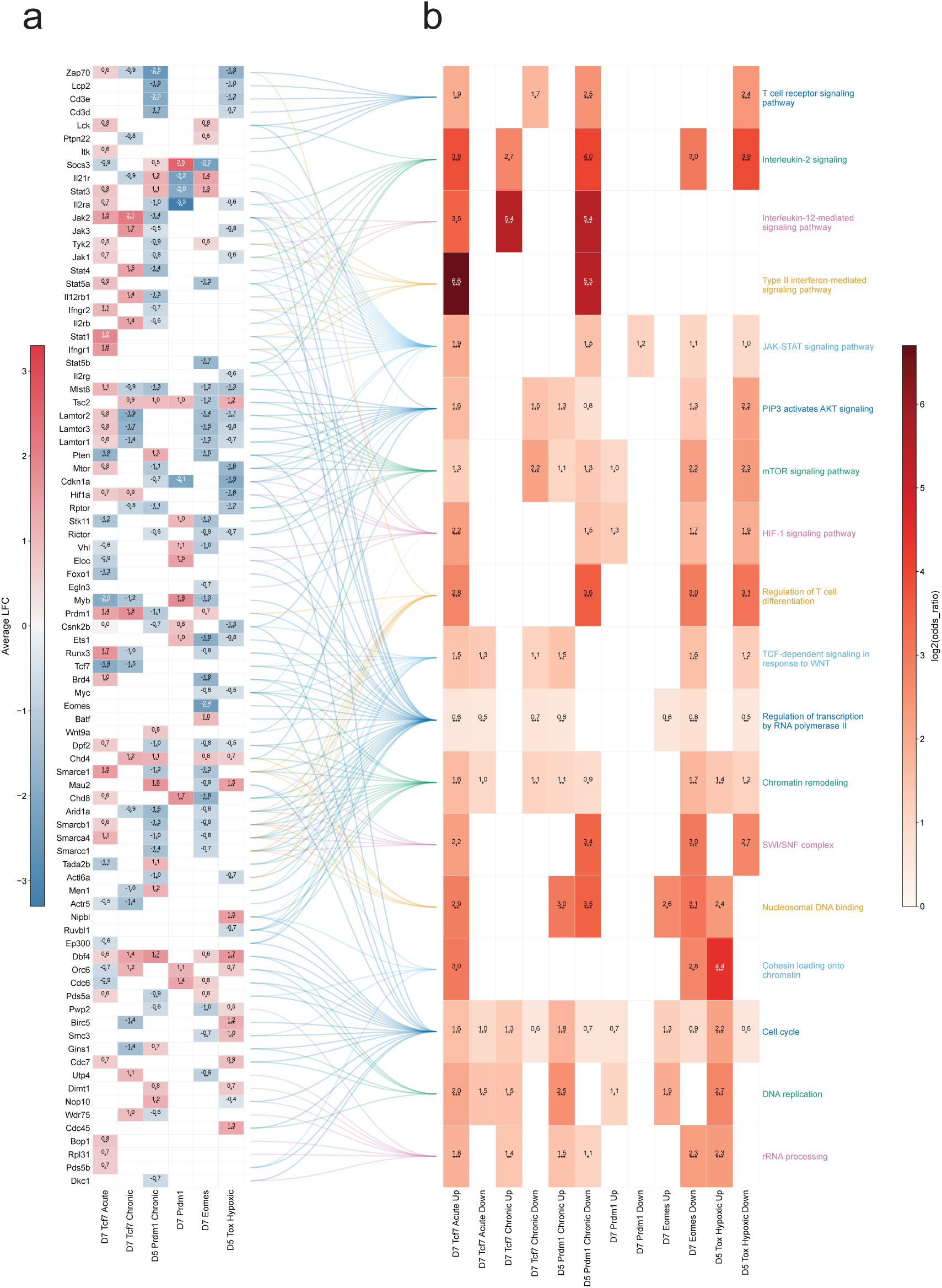
Pathway enrichment analysis for CRISPR screen results. (a) Gene-level heatmap of manually curated *in vitro* screen hits linked to the pathway terms in (b) across six condition-specific genome-wide CRISPR screen comparisons. Values are PoolQ Average LFC across guides and are shown for screen-gene pairs with nominal gene-level P <= 0.05. Positive values indicate enrichment in the high-marker bin, and negative values indicate enrichment in the low-marker bin (Table S1.5). Significance labels: ‘***’, P < 0.001; ‘**’, P < 0.01; ‘*’, P <= 0.05. (b) Pathway enrichment heatmap for the terms linked to genes in (a). For each screen comparison, genes with nominal gene-level P <= 0.05 were split by the sign of PoolQ Average LFC: positive values indicate enrichment in the high-marker bin, and negative values indicate enrichment in the low-marker bin. Each gene set was tested for enrichment in selected GO Biological Process, GO Cellular Component, GO Molecular Function, Reactome, KEGG Mouse, and WikiPathways Mouse terms using one-sided Fisher’s exact tests. Values are log2(odds ratio) and are shown for pathway-direction pairs with P < 0.05.

**Figure S3.**
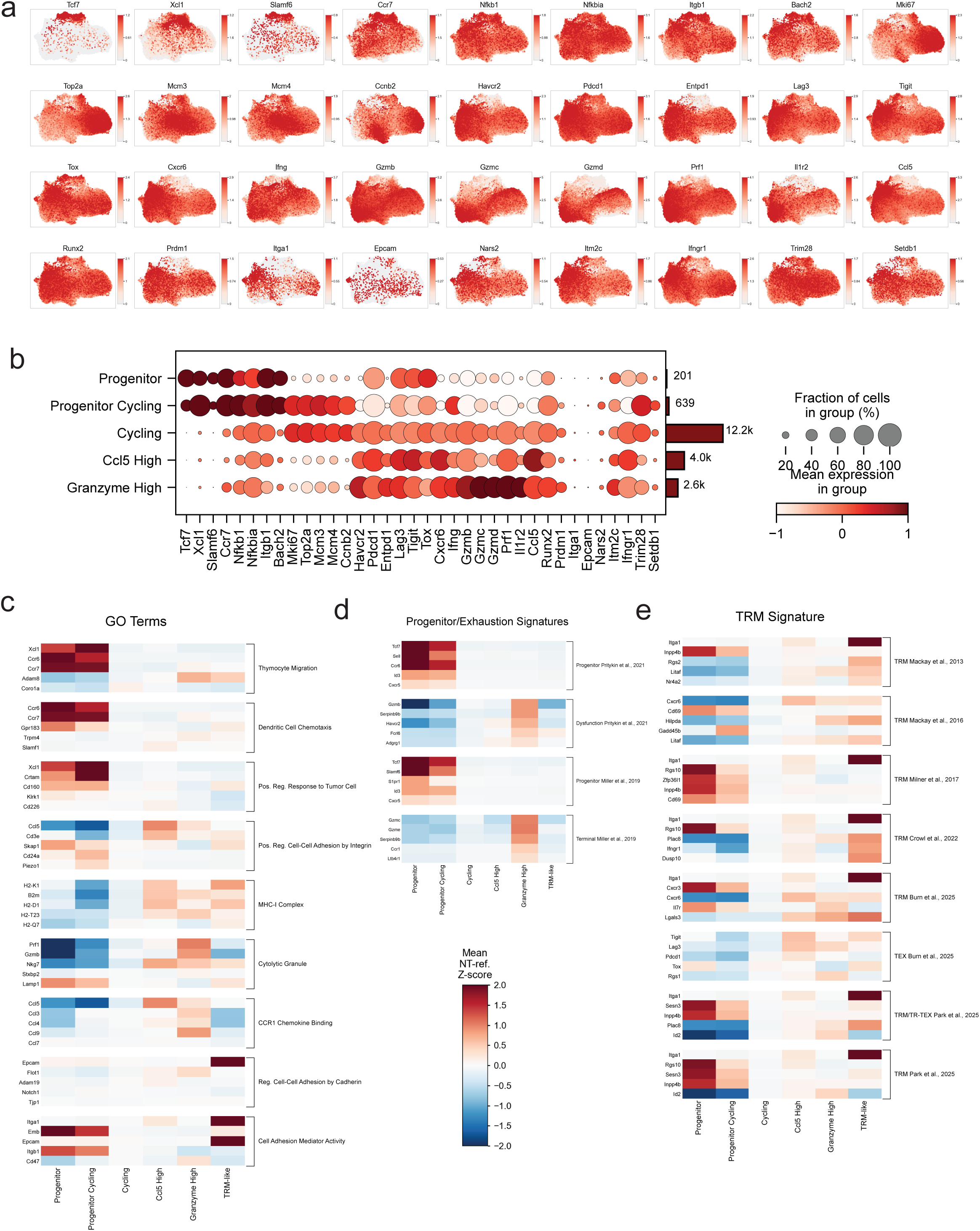
Additional Perturb-seq analysis. (a) UMAP plots showing library-size-normalized expression of key marker genes in the 10-gene Perturb-seq dataset. (b) Same as Figure 2c, but restricted to NT cells and excluding the TRM-like state. (c) Gene-level heatmaps showing representative genes underlying the GO term-derived signature programs in Figure 2e. Columns are the six cell states and rows are genes grouped by program. Heatmap values are the mean NT-referenced log-normalized z-scored expression. We manually selected 5 genes for each program based on manual curation and large differences across cell states. (d) Gene-level heatmaps showing representative genes underlying the progenitor and dysfunction programs used in Figure 2e^2,60^. The same manual curation and plotting procedure described in (c) was applied to these signatures. (e) Gene-level heatmaps showing representative genes underlying the TRM- and TEX-related programs used in Figure 2e. The same manual curation and plotting procedure described in (c) was applied to these signatures.

**Figure S4.**
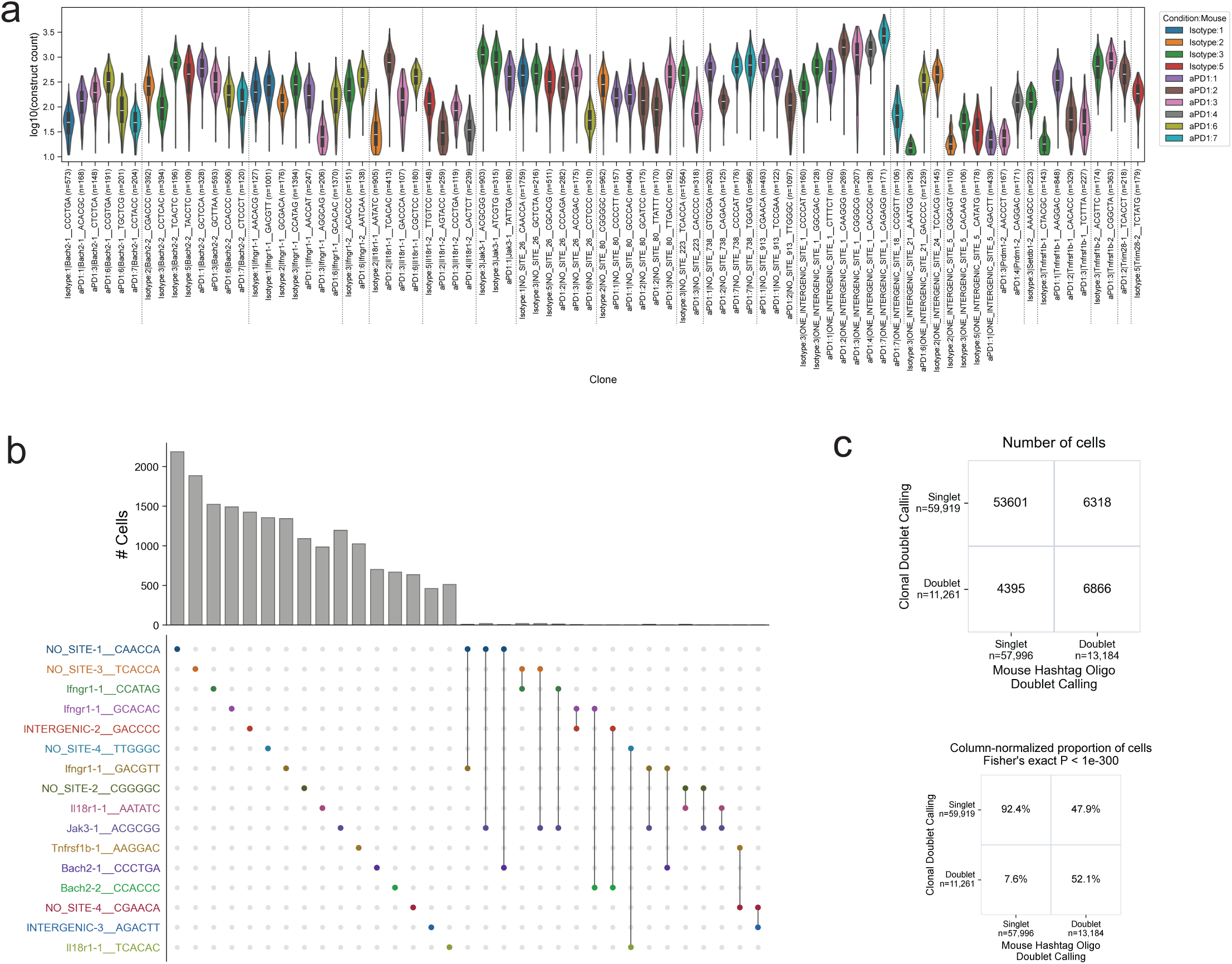
Clonal assignment in barcoded Perturb-seq data. (a) Distribution of construct UMI counts across cells within single-construct clones with more than 100 cells in the 10-gene Perturb-seq dataset. Each violin shows the distribution of log10(construct count) across cells within a clone, where each construct is defined by guide RNA, two underscores, and a 6-bp clonal barcode. X-axis labels report condition:mouse, construct identity, and the number of cells in the clone (‘n’). Clones are ordered by guide, with Isotype mice shown before anti-PD-1 mice for each guide. Dotted lines separate guides. (b) UpSet plot of putative clone construct combinations in the 10-gene Perturb-seq dataset. Columns show selected putative clones. Top bars show the number of cells assigned to each displayed putative clone, and filled circles denote the constituent constructs. To select the putative clones for display, we first selected the 20 clones with the most cells and collected the constructs carried by those clones. We then considered putative clones defined before the linear-program doublet filter from construct calls with count > 10, restricted to construct combinations observed in at least 3 cells. Finally, to simplify the display, we retained only construct combinations involving selected constructs that were observed together in a multi-construct putative clone. (c) 2×2 tables comparing singlet and doublet assignments from clonal doublet calling and mouse hashtag oligo doublet calling in the 10-gene Perturb-seq dataset. The count table reports cell counts. The column-normalized table reports the proportion of cells within each mouse hashtag oligo category assigned as clonal singlets or doublets. The p-value is from a one-sided Fisher’s exact test for positive association between clonal doublet calling and mouse hashtag oligo doublet calling.

**Figure S5.**
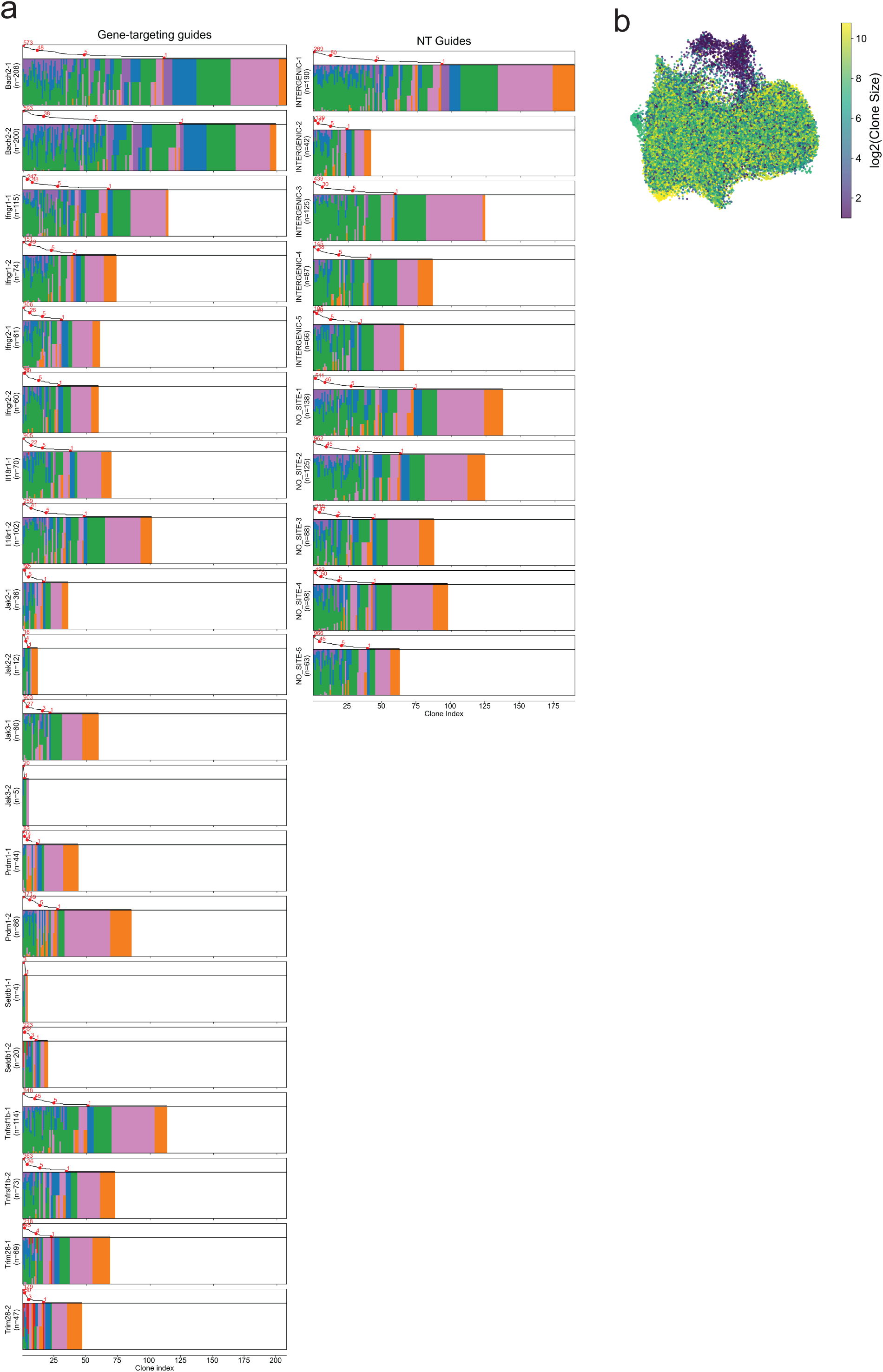
Additional clonal analysis of Perturb-seq data. (a) Same as Figure 3b, but showing clone-size and cell-state heterogeneity separately for each guide. (b) UMAP plot colored by clone size.

**Figure S6.**
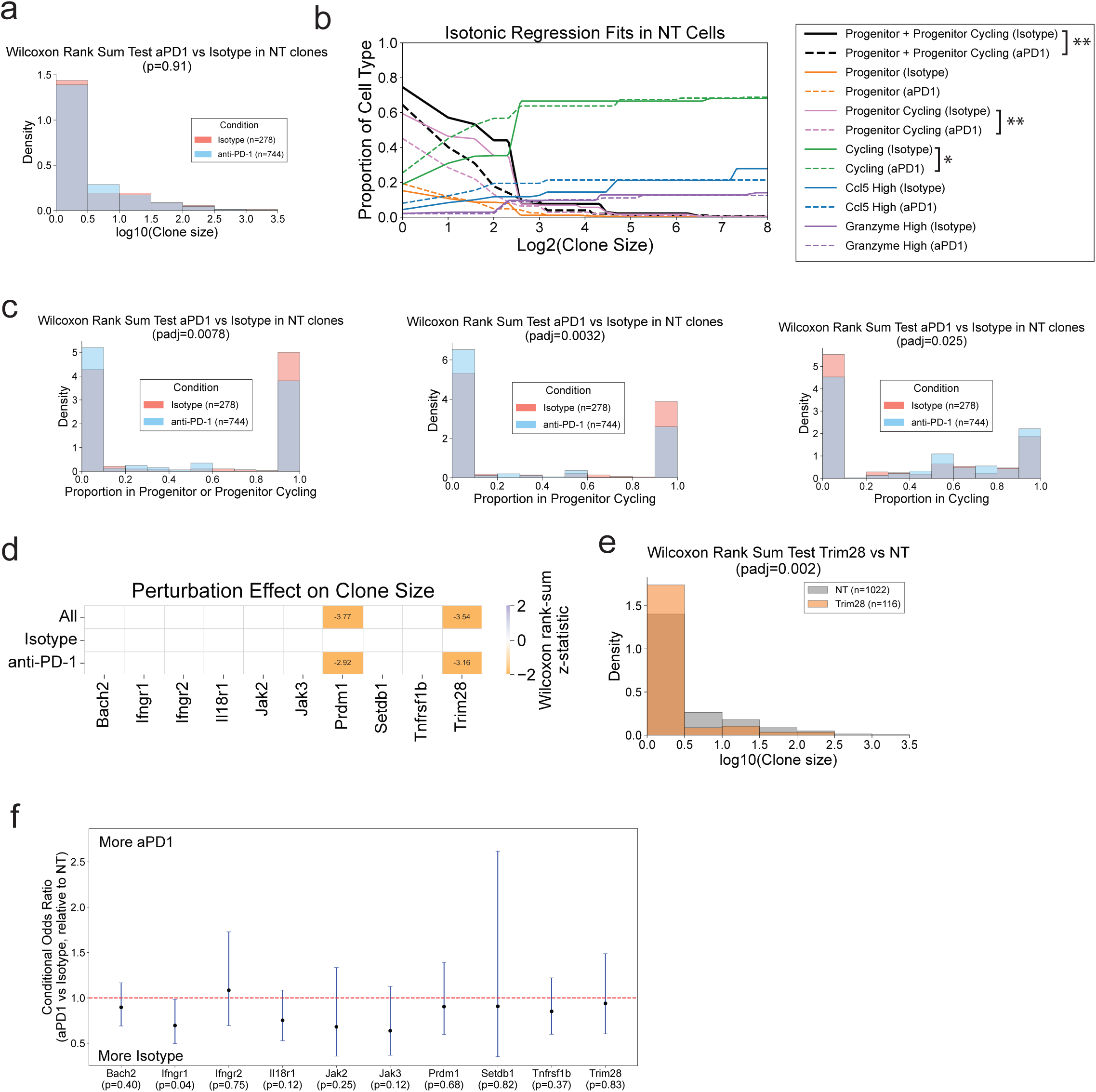
Perturbation effect on clonal size. (a) Overlapping histograms of NT clone sizes for Isotype and anti-PD-1 conditions. The x-axis shows log10(clone size) and the y-axis shows density. Here, ‘n’ denotes the number of NT clones. (b) Isotonic regression fits for NT clones split by Isotype and anti-PD-1, showing the relationship between log2(clone size) and the proportion of cells in each state. The black line denotes the combined Progenitor plus Progenitor Cycling fraction. Significance between anti-PD-1 and Isotype NT clone-level cell-type proportions was assessed with two-sided Wilcoxon rank-sum tests and Benjamini-Hochberg correction across the displayed states. Significance labels: ‘**’, adjusted P < 0.01; ‘*’, adjusted P < 0.05. (c) Overlapping histograms of NT clone-level cell-state proportions for Isotype and anti-PD-1 conditions. (d) Heatmap showing perturbation effects on clone size. Rows show all cells pooled, Isotype-treated cells, and anti-PD-1-treated cells. Using a Wilcoxon rank-sum test, log2(clone size) for perturbed clones was compared with NT controls. The effect size is shown when BH-adjusted P < 0.05. (e) Overlapping histograms of log10 clone size for NT and Trim28. This panel shows the Trim28 example underlying the pooled comparison in (d). (f) Forest plot of conditional odds ratios comparing the anti-PD-1-versus-Isotype clone-count ratio for each perturbation against the corresponding NT ratio in the 10-gene Perturb-seq dataset. Blue bars denote exact conditional 95% confidence intervals from 2×2 contingency tables with rows NT versus perturbation and columns Isotype versus anti-PD-1. X-axis labels report two-sided Fisher’s exact-test P values.

**Figure S7.**
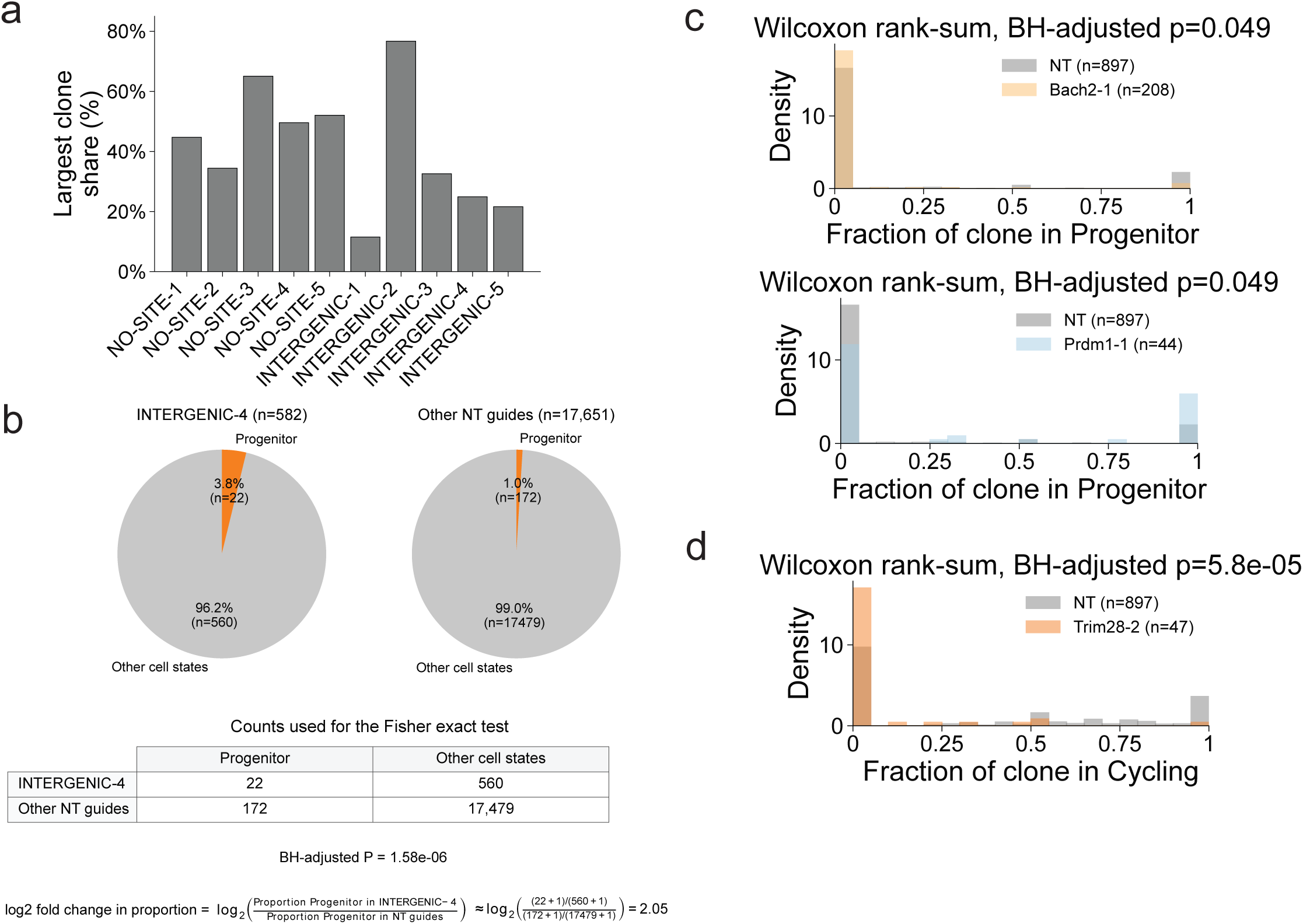
Additional analysis of clonal distributions. (a) Bar plot across control guides showing the percentage of guide-assigned cells contributed by the largest recovered clone for that guide in the 10-gene Perturb-seq dataset. (b) Example control-versus-control comparison underlying the Figure 4c cell-based Fisher’s test for INTERGENIC-4 in the Progenitor state. Pie charts show the fraction of Progenitor versus all other cell states for INTERGENIC-4 cells and for the remaining NT guides. The 2×2 contingency table reports the corresponding cell counts used for the Fisher’s exact test. Benjamini-Hochberg-adjusted P values are calculated across all guides for the Progenitor state. The log2 odds ratio was computed with a +1 pseudocount and was used as the value plotted in Figure 4c. (c) Histograms showing the distribution across clones of the fraction of cells annotated as Progenitor within each clone. These histograms underlie the comparison made in Figure 4d for the Progenitor state. The top histogram compares NT clones against Bach2-1 clones, and the bottom histogram compares NT clones against Prdm1-1 clones; when multiple guides for a perturbation were significant in Figure 4d, the guide with the larger absolute Wilcoxon rank-sum z-statistic was shown. Significance was assessed with two-sided Wilcoxon rank-sum tests and Benjamini-Hochberg correction within the cell state across guides. (d) Same as (c), but showing clone-level Cycling fractions for NT versus Trim28-2 clones, corresponding to the depletion of Cycling cells detected in Figure 4d.

**Figure S8.**
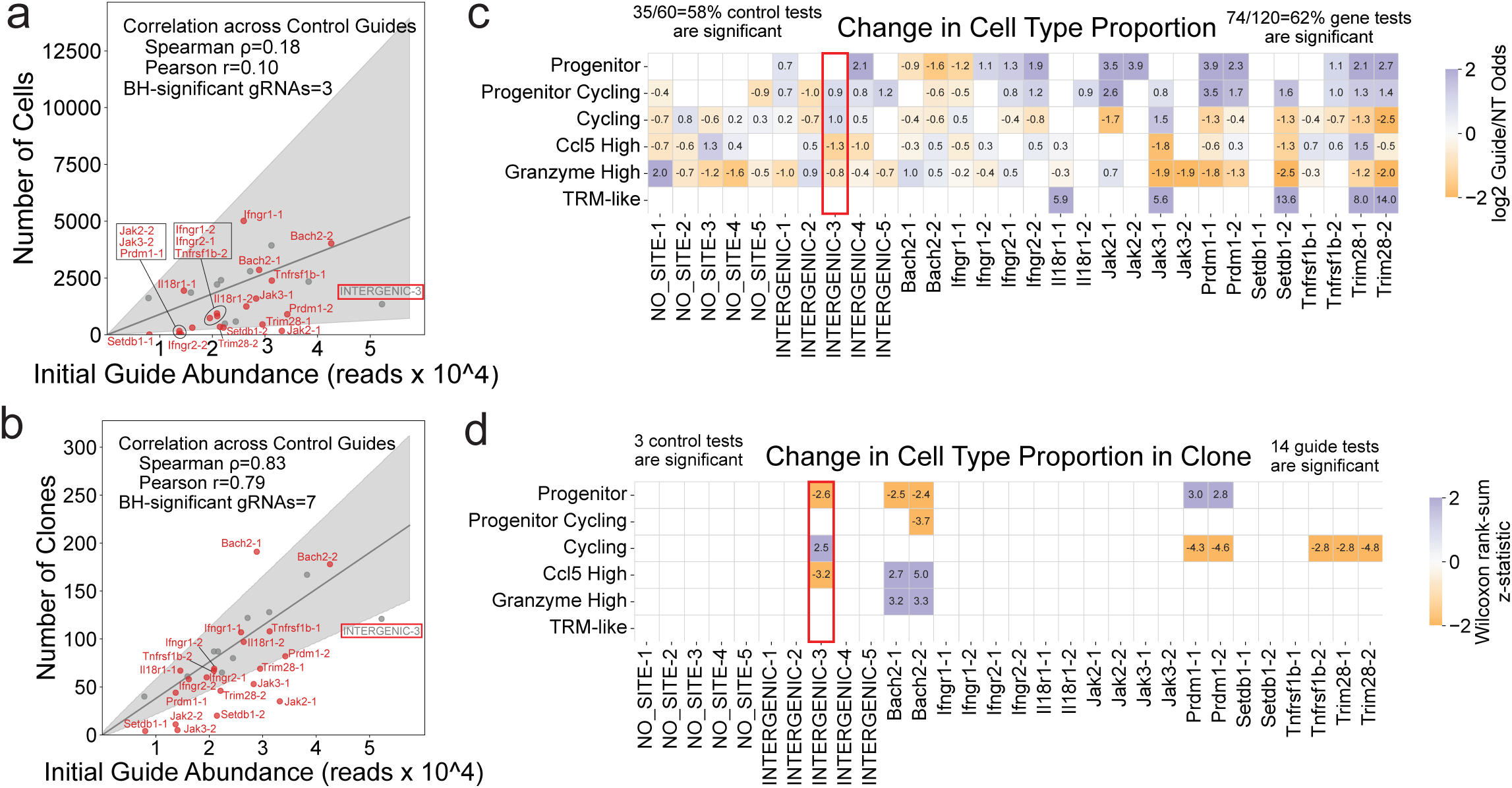
Cell-based and clonal statistical testing. (a-d) Same as Figure 4a-d, but including all guides; this includes INTERGENIC-3, the intergenic control gRNA excluded from Figure 4a-d and highlighted in red.

**Figure S9.**
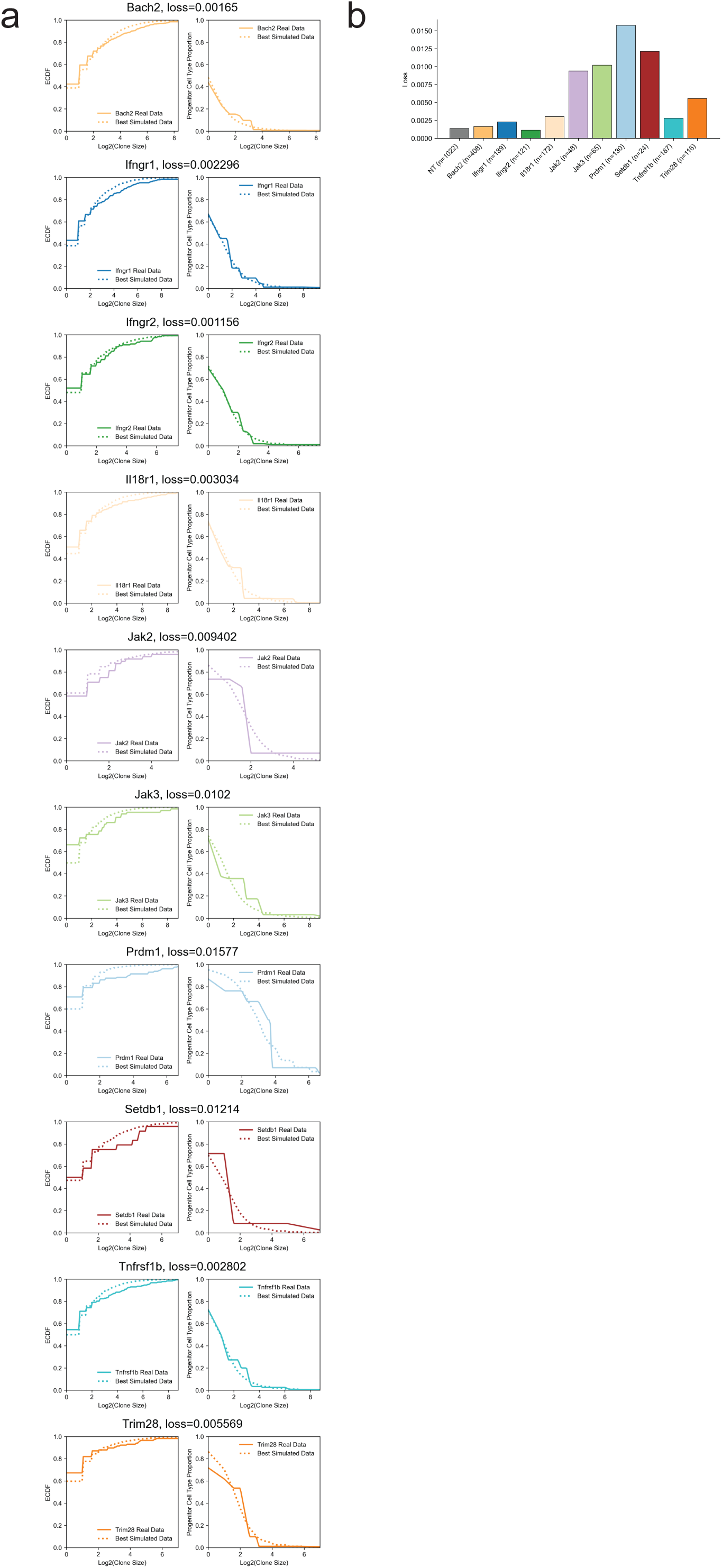
Clonal modeling of cell differentiation. (a) Same summary statistics as Figure 4g, but shown for the best-loss parameter set for each perturbation. Left, ECDF of log2(clone size) for the perturbation’s real data and best simulated data. Right, progenitor cell type proportion versus log2(clone size) for the perturbation’s real and best simulated clones, with curves fit by isotonic regression. (b) Bar plot showing the loss for each modeled perturbation (n is the number of clones).

**Figure S10.**
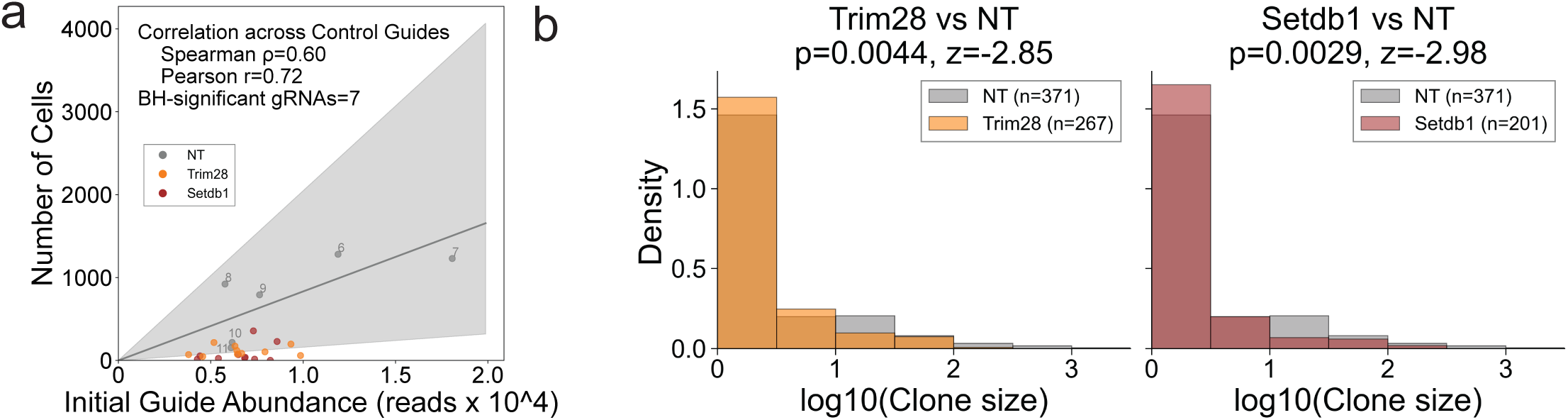
Clonal size distribution in the Trim28/Setdb1 Perturb-seq experiment. (a) Same as Figure 5b, but using recovered cell counts rather than recovered clone counts. (b) Overlapping histograms of log10 clone size for NT versus Trim28 (left) and NT versus Setdb1 (right) after excluding lymph-node cells. The y-axis shows density. Two-sided Wilcoxon rank-sum P values and z-statistics are shown for each pooled comparison.

**Figure S11.**
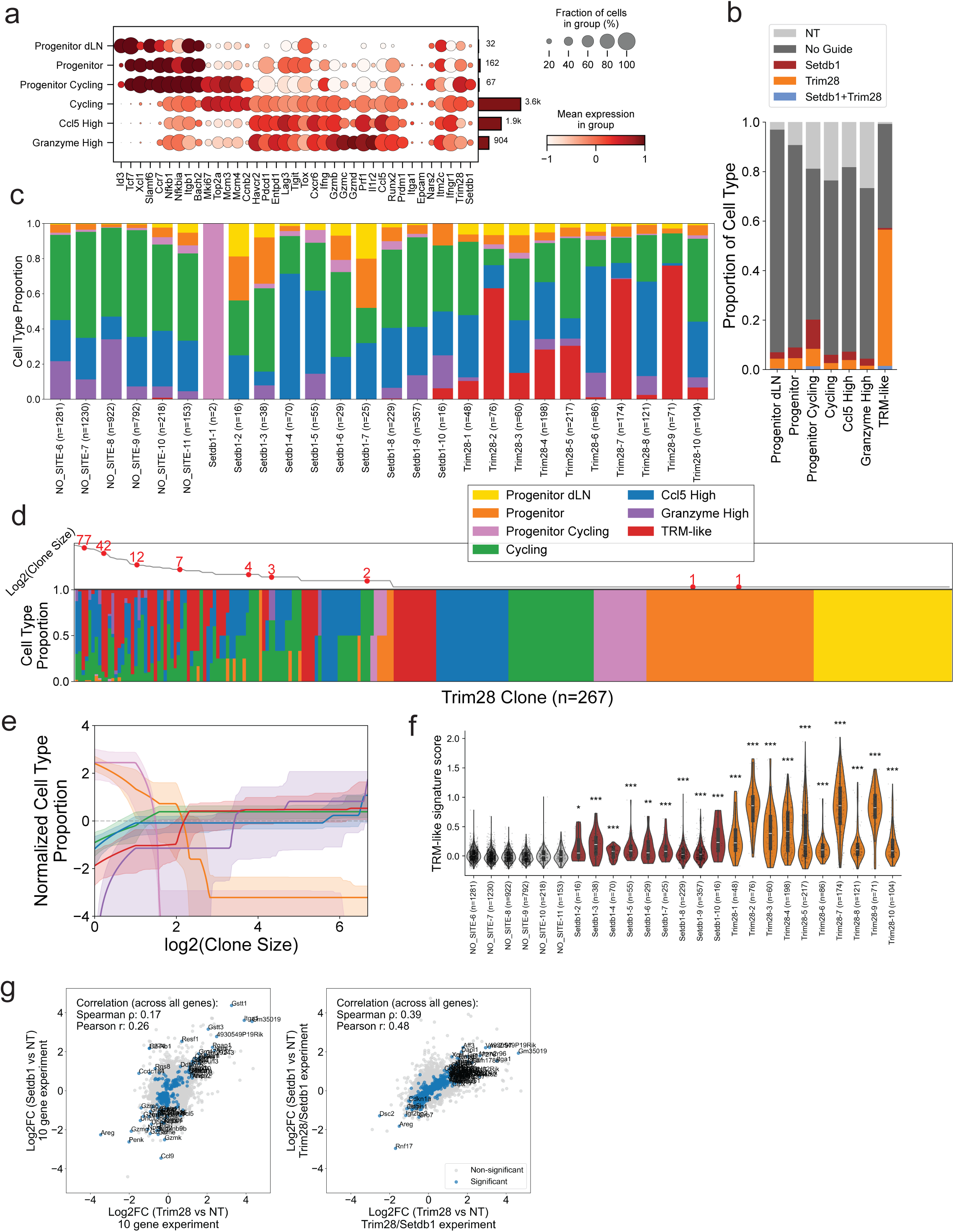
Additional analysis of the Trim28/Setdb1 Perturb-seq experiment. (a) Dot plot visualizing expression of key marker genes across NT cell states in the Trim28/Setdb1 Perturb-seq dataset, excluding the TRM-like state. Dot size indicates the fraction of cells expressing each marker, color indicates mean expression, and the bar plot at right shows the number of cells per state. (b) Stacked bar chart showing the relative abundance of perturbation categories within each cell state, as in Figure 5g, but including cells with no detected guide as a separate ‘No Guidè category shown in dark gray. (c) Stacked bar chart showing guide-level cell-state proportions in the Trim28/Setdb1 Perturb-seq dataset. ‘n’ denotes the number of cells. (d) Clone-level cell-state composition for Trim28 clones from the Trim28/Setdb1 Perturb-seq dataset, analogous to Figure 3b. Clones are ordered by log2(clone size), then by cell-type composition, along the x-axis. The top panel shows log2(clone size) across clones; red dots mark clones selected by uniform sampling weighted by log2(clone size), and labels report their clone sizes. The bottom panel shows stacked cell-type proportions for each clone. (e) Clone-size-normalized cell-state enrichment analysis for Trim28 clones from the Trim28/Setdb1 Perturb-seq dataset after excluding lymph-node cells, analogous to Figure 3f. (f) Violin plot showing guide-level TRM-like signature scores in the Trim28/Setdb1 Perturb-seq dataset using the TRM-like signature genes (Table S5.2). Guides with more than 10 cells are shown. Significance was assessed with one-sided Mann-Whitney U tests comparing each guide against pooled NT cells; x-axis labels report the number of cells (‘n’). Significance labels: ‘***’, P < 0.001; ‘**’, P < 0.01; ‘*’, P < 0.05. (g) Scatter plots comparing Trim28-versus-NT log2 fold changes (x-axis) and Setdb1-versus-NT log2 fold changes (y-axis) within each experiment. Left, the 10-gene Perturb-seq experiment. Right, the Trim28/Setdb1 Perturb-seq experiment. Points denote shared genes; blue points were significant in both perturbation comparisons within the displayed experiment, and gray points were not.

**Figure S12.**
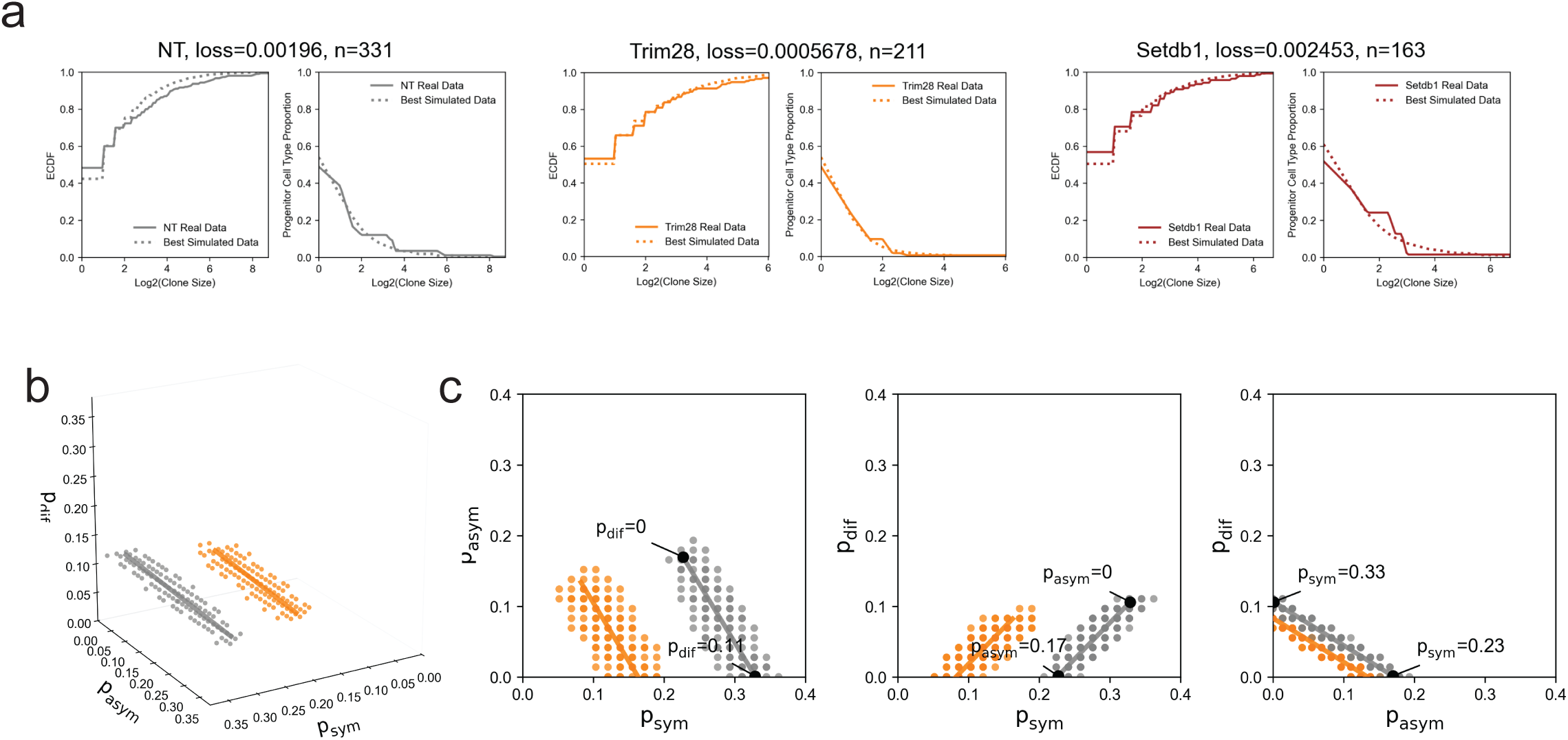
Clonal modeling for the Trim28/Setdb1 Perturb-seq data. (a) Same summary statistics as Figure 4g, but shown for the best-loss parameter set for NT, Trim28, and Setdb1 in the Trim28/Setdb1 Perturb-seq dataset. ‘n’ is the number of clones. (b) Same as Figure 4f, but showing the 100 lowest-loss parameter sets for NT and Trim28 in the Trim28/Setdb1 Perturb-seq dataset. (c) Same as Figure 4h, but for the NT and Trim28 lowest-loss parameter sets in the Trim28/Setdb1 Perturb-seq dataset.

**Figure S13.**
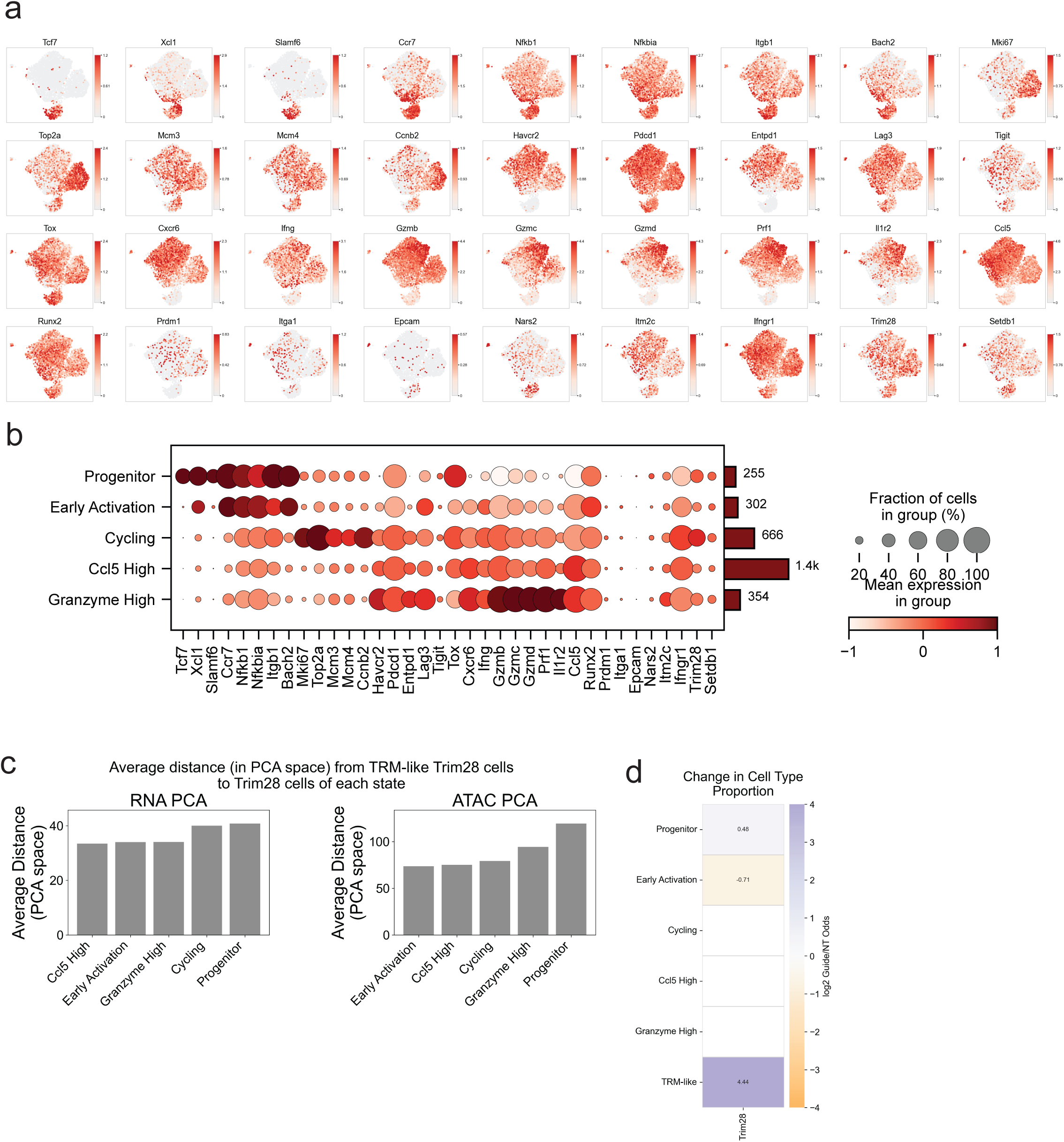
Additional analysis of the multi-omic data. (a) UMAP plots showing library-size-normalized expression of key marker genes in the multiome RNA dataset. (b) Same as Figure 6b, but restricted to control cells and excluding the TRM-like state. (c) Same analysis as Figure 5m, but applied to the multiome Trim28 cells. Left, mean Euclidean distances computed in RNA PCA space (first 30 PCs). Right, mean Euclidean distances computed in ATAC PCA space (first 40 PCs). (d) Same analysis as Figure 4c, but comparing Trim28 versus Control cells in the multiome RNA dataset. Values are shown when BH-adjusted two-sided P < 0.05 across the cell states.

**Figure S14.**
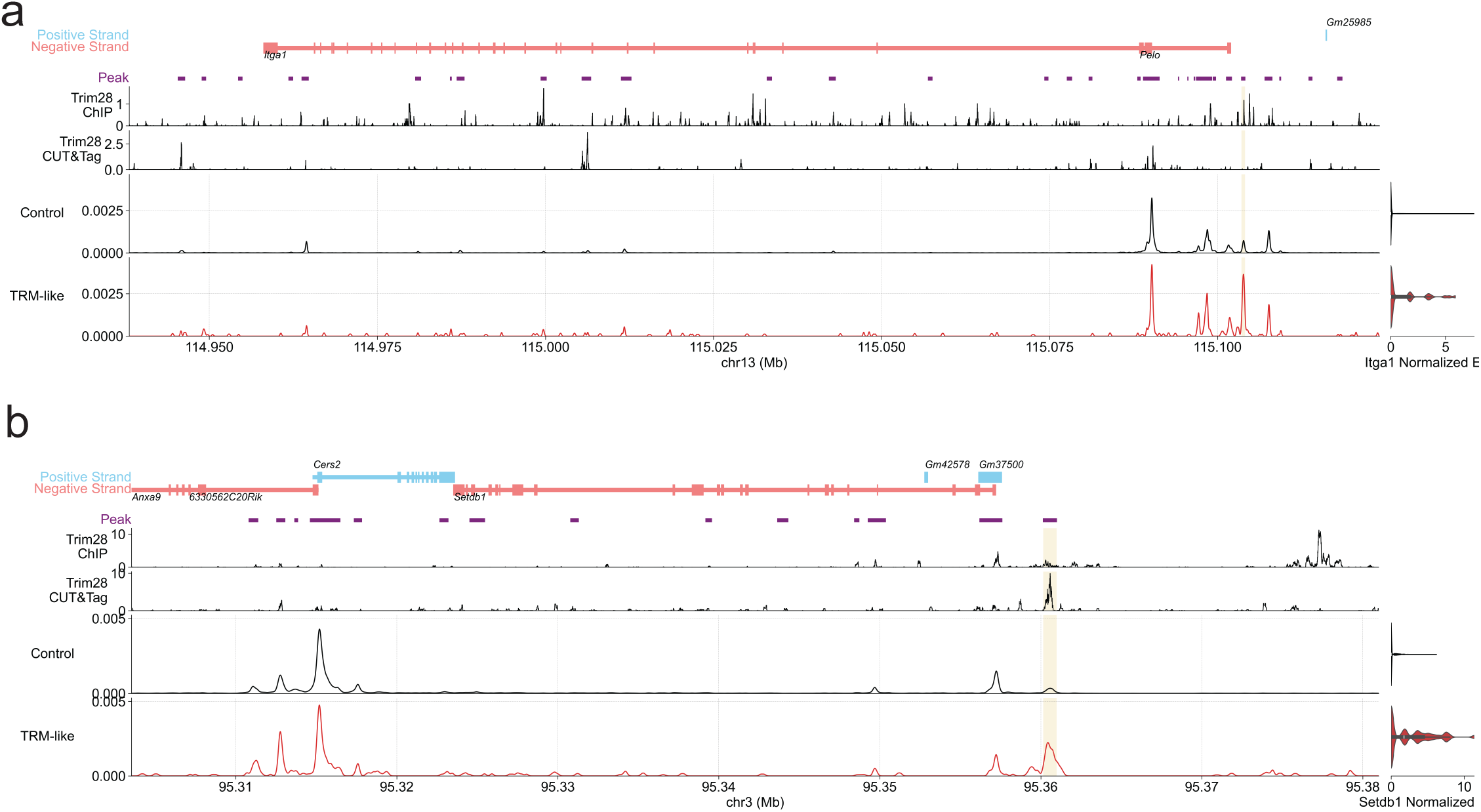
Genomic tracks for *Itga1* and *Setdb1*. (a) Same layout as Figure 6e, but showing the *Itga1* locus. Control and TRM-like cells are shown, and the shaded region marks the *Itga1*-assigned ATAC peak that was significant in the Figure 6d comparison of Trim28 TRM-like cells versus non-TRM-like Control cells. The right panel shows the corresponding normalized scRNA-seq expression of *Itga1*. (b) Same layout as Figure 6e, but showing the *Setdb1* locus. Control and TRM-like cells are shown, and the shaded region marks the *Setdb1*-assigned ATAC peak that was significant in the Figure 6d comparison of Trim28 TRM-like cells versus non-TRM-like Control cells. The right panel shows the corresponding normalized scRNA-seq expression of *Setdb1*.

**Figure S15.**
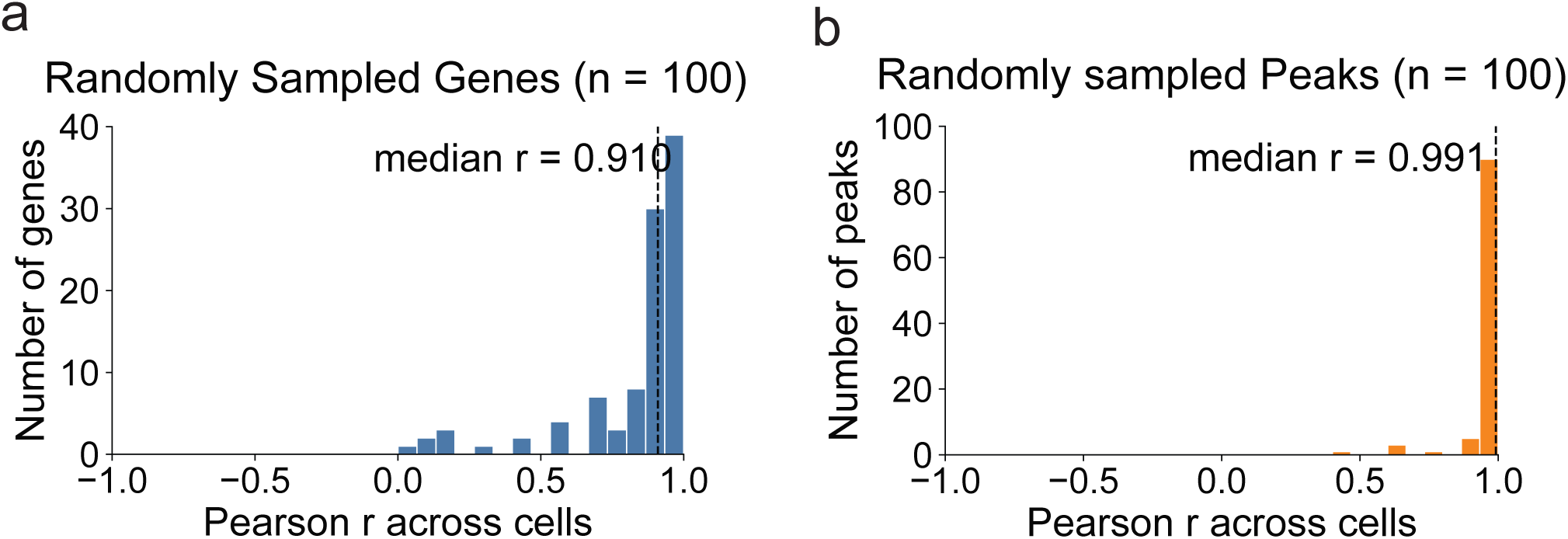
Concordance of the multiome preprocessing pipelines. (a) Histogram showing the distribution of Pearson correlations across cells for 100 randomly sampled retained genes in the scRNA-seq component of the multiome dataset. For each gene, counts reconstructed from the raw multiome scRNA-seq BAM files were compared with counts derived from Cell Ranger. (b) Histogram showing the distribution of Pearson correlations across cells for 100 randomly sampled retained peaks in the scATAC-seq component of the multiome dataset. For each peak, insertion counts reconstructed from the raw multiome scATAC-seq BAM files were compared with peak counts derived from Cell Ranger.

